# Alzheimer’s disease linked Aβ42 exerts product feedback inhibition on γ-secretase impairing downstream cell signaling

**DOI:** 10.1101/2023.08.02.551596

**Authors:** Katarzyna Marta Zoltowska, Utpal Das, Sam Lismont, Thomas Enzlein, Masato Maesako, Mei CQ Houser, María Luisa Franco, Burcu Özcan, Diana Gomes Moreira, Dmitry Karachentsev, Ann Becker, Carsten Hopf, Marçal Vilar, Oksana Berezovska, William Mobley, Lucía Chávez-Gutiérrez

**Author notes:** **Corresponding authors:** Lucía Chávez-Gutiérrez VIB-KU Leuven Center for Brain & Disease Research Herestraat 49 bus 602 3000 Leuven, Belgium, Mobley William University of California, San Diego School of Medicine George Palade Labs (GPL) 359 9500 Gilman Drive La Jolla, CA 92037-0662.

## Abstract

Amyloid β (Aβ) peptides accumulating in the brain are proposed to trigger Alzheimer’s disease (AD). However, molecular cascades underlying their toxicity are poorly defined.

Here, we explored a novel hypothesis for Aβ42 toxicity that arises from its proven affinity for γ-secretases. We hypothesized that the reported increases in Aβ42, particularly in the endolysosomal compartment, promote the establishment of a product feedback inhibitory mechanism on γ-secretases, and thereby impair downstream signaling events.

We show that human Aβ42 peptides, but neither murine Aβ42 nor human Aβ17-42 (p3), inhibit γ-secretases and trigger accumulation of unprocessed substrates in neurons, including C-terminal fragments (CTFs) of APP, p75 and pan-cadherin. Moreover, Aβ42 treatment dysregulated cellular homeostasis, as shown by the induction of p75-dependent neuronal death in two distinct cellular systems.

Our findings raise the possibility that pathological elevations in Aβ42 contribute to cellular toxicity via the γ-secretase inhibition, and provide a novel conceptual framework to address Aβ toxicity in the context of γ-secretase-dependent homeostatic signaling.

## INTRODUCTION

Γ-secretases are ubiquitously expressed intramembrane proteases best known for their pathogenic roles in Alzheimer’s disease (AD) (*1*). Aberrant processing of the amyloid precursor protein (APP) by γ-secretases leads to the production of longer, aggregation-prone amyloid β (Aβ) peptides that contribute to neurodegeneration (*2*).

In addition, γ-secretases process many other membrane proteins, including NOTCH, ERB-B2 receptor tyrosine kinase 4 (ERBB4), N-cadherin (NCAD) and p75 neurotrophin receptor (p75-NTR) (*3, 4*). The processing of multiple substrates links their activity to a broad range of downstream signaling pathways (*5, 6*), including those critical for neuronal function. It is noteworthy that treatment with γ-secretase inhibitors caused cognitive worsening in AD patients (*7*), while full genetic inhibition of these enzymes in the adult mouse brain led to neurodegenerative phenotypes (*8–12*). The underlying mechanisms by which the deficits in γ-secretase activity impair neuronal function are yet to be defined.

Γ-secretase activity is exerted by a family of highly homologous multimeric proteases composed of presenilin (PSEN1 or PSEN2), nicastrin (NCSTN), anterior pharynx defective 1 (APH1A or B) and presenilin enhancer 2 (PEN2) subunits. The proteolytic activities of these complexes are promoted by the low pH of the endosomal and lysosomal compartments, wherein the amyloidogenic processing of APP occurs (*13*). In the amyloidogenic pathway the proteolytic processing of APP by β-secretase (BACE) releases a soluble APP ectodomain and generates a membrane-bound C-terminal fragment (β-CTF or APP_C99_) (*14*). APP_C99_ is then sequentially processed within the membrane by γ-secretase complexes **(Figure 1A)** (*15–19*). An initial endopeptidase (ε-) cut releases APP intracellular domain (AICD) into the cytosol and generates a *de novo* substrate (either Aβ49 or Aβ48 peptide) that undergoes successive γ-cleavages until a shortened Aβ peptide can be released into the luminal or extracellular environment. The efficiency of the sequential cleavage mechanism (i.e. processivity) determines the length of Aβ (37-43 amino acid long peptides), which in turn influences the aggregation and neurotoxic properties of the peptides produced (*2, 20, 21*). In the non-amyloidogenic pathway APP is cleaved by α- and γ-secretases to generate a spectrum of p3 peptides, which lack the first 1-16 amino acids of Aβ **(Figure 1A)**. Despite their relatively high hydrophobicity and aggregation prone behavior, the p3 peptides are not linked to AD pathogenesis (*22–24*). In fact, mutations that promote the amyloidogenic processing of APP are associated with AD (*25, 26*), whereas those that favor the alternative, non-amyloidogenic pathway protect against the disease (*24, 27*).

**Figure 1.**
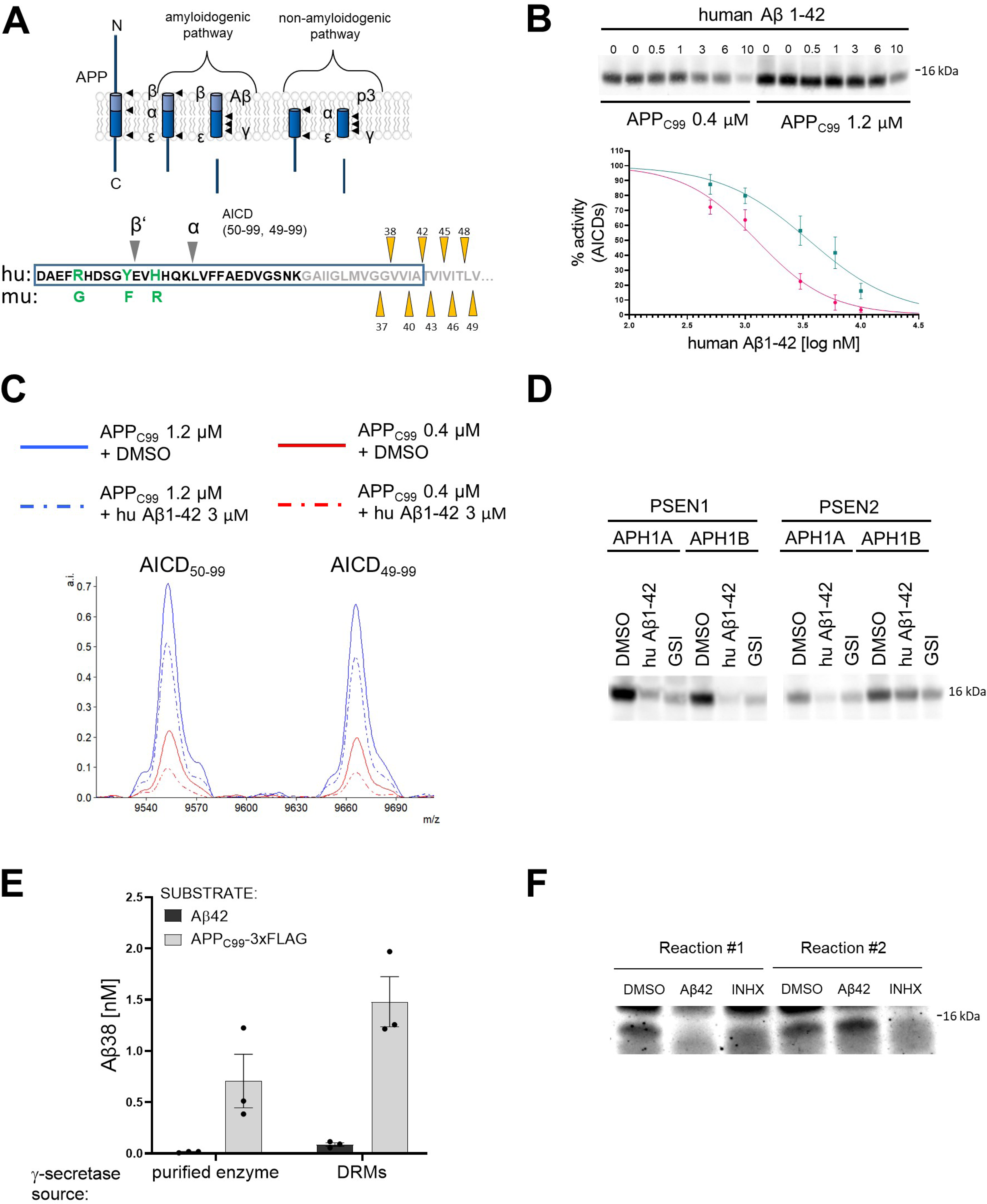
Human Aβ42 peptide inhibits γ-secretase-mediated proteolysis of APP_C99_. (A) The scheme depicts the γ-secretase-mediated cleavage of APP, leading to the generation of Aβ and p3 peptides. The N-terminal sequence of APP_C99_ /Aβ is shown in the lower panel. The differences in the amino acid sequence of human (hu) vs murine (mu) Aβ peptides and the positions of β’- and α-cleavages (that precede the generation of Aβ11-42 and p3 17-42 peptides, respectively) are indicated. The transmembrane domain is labelled in grey and the sequence of Aβ42 is presented within a rectangle. The initial γ-secretase endopeptidase cut may occur at one of two different positions on APP, generating two different *de novo* substrates that are further processed by carboxypeptidase-like cleavages as follows: Aβ49→Aβ46→Aβ43→Aβ40→Aβ37 or Aβ48 →Aβ45→Aβ42→Aβ38. Aβ38, Aβ40 and Aβ42 peptides are the major products under physiological conditions. The triangles mark the sequential cleavage positions. (B) The western blot presents AICD products generated *de novo* in detergent-based γ-secretase activity assays using APP_C99_-3xFLAG at 0.4 μM or 1.2 μM as substrate. To test the inhibitory properties of human Aβ1-42, the peptide was added to the activity assays at concentrations ranging from 0.5 to 10 μM. DMSO at 2.5% was used as a vehicle control. The graphs present the quantification of the western blot bands corresponding to AICDs. The pink and green lines correspond to 0.4 μM and 1.2 μM substrate concentrations, respectively. The data are normalized to the AICD levels generated in the DMSO conditions, considered as 100%. The data are presented as mean ± SEM, N=6-16. (C) Mass spectrometry-based analysis of *de novo* generated AICD levels in *in vitro* cell-free γ-secretase activity assays containing 3 μM human Aβ1-42 or vehicle (0.6% DMSO) indicate that human Aβ1-42 inhibits both product lines. (D) The western blot presents *de novo* generated AICDs in detergent-based γ-secretase activity assays using 0.4 μM APP_C99_-3xFLAG as substrate and wild type γ-secretases composed of different PSEN (1 or 2) and APH1 (A or B) subunits in the presence of vehicle, human Aβ1-42 at 3 μM or GSI (InhX) at 10 μM concentration. (E) The graph presents ELISA quantification of Aβ1-38 peptides generated in detergent- or detergent resistant membranes (DRM)-based γ-secretase activity assays using either human Aβ1-42 at 10 μM or APP_C99_-3xFLAG at 1.5 μM as substrates. The data are presented as mean ± SEM, N=3. (F) The western blot presents *de novo* generated AICDs in detergent-based γ-secretase activity assays using 0.4 μM APP_C99_-3xFLAG as a substrate and γ-secretase immobilized on sepharose beads. During the first round, the activity assay was supplemented with 3 μM Aβ1-42, 10 μM GSI or DMSO vehicle. After this first round, the γ-secretase-beads were washed to remove the peptide and inhibitor, respectively, fresh substrate was added, and the reaction proceeded for the second round. The analysis demonstrates the reversibility of the Aβ1-42-mediated inhibition.

In familial AD (FAD), mutations in PSENs or in APP/Aβ (i.e. changes effecting both the enzyme and its substrate) promote the generation of longer Aβ peptides, including Aβ42, but also Aβ43 (*28–33*). These peptides accumulate in the brain and are a hallmark of AD, in both familial and sporadic forms. In the latter (SAD), the accumulation of longer, aggregation prone peptides results from their inefficient clearance (*34, 35*). Irrespective of the mechanism, the accumulation of Aβ in the brain begins decades before the onset of clinical symptoms and is proposed to trigger toxic cascades via poorly understood mechanisms (*2*).

Aggregation of Aβ42 into soluble toxic oligomers has been proposed as a prerequisite for its toxicity (*36–39*). However, given the broad spectrum of brain-derived Aβ peptides and their assemblies, it seems plausible that Aβ contribute via a number of mechanisms to AD pathogenesis. Of note, increasing evidence suggests that in addition to Aβ (*2*), its precursor, the 99 amino acid C-terminal fragment (i.e. APP_C99_) (*40*) also mediates pathogenic mechanisms (*41–43*).

Herein, we explored a novel hypothesis for Aβ42 toxicity that arises from its proven affinity for γ-secretases. We hypothesized that pathological increases in Aβ levels in the AD brain lead to the establishment of a product feedback inhibitory mechanism, wherein Aβ1-42 competes with membrane-associated substrates for γ-secretase processing. This hypothesis is supported by our reported observations that Aβs with low affinity for γ-secretase, when present at relatively high concentrations, can compete with the longer, higher affinity APP_C99_ substrate for binding and processing by the enzyme (*30*). This hypothetical inhibitory feedback would result in the accumulation of unprocessed γ-secretase substrates and reduced levels of their soluble intracellular domain (ICD) products, possibly disrupting downstream signaling cascades.

To test the hypothesis, we investigated whether Aβ peptides can exert inhibition on γ-secretase. Our kinetic analyses demonstrate that human Aβ1-42 inhibits γ-secretase-mediated processing of APP_C99_ and other substrates. Strikingly, neither murine Aβ1-42 nor human p3 (17-42 amino acids in Aβ) peptides exerted inhibition under similar conditions. We also show that human Aβ1-42-mediated inhibition of γ-secretase activity results in the accumulation of unprocessed CTFs of APP, p75 and pan-cadherins. To evaluate the impact of the Aβ-driven inhibition on cellular signaling, we analyzed p75-dependent activation of caspase 3 in basal forebrain cholinergic neurons (BFCNs) and PC12 cells. These analyses demonstrate that, as seen for γ-secretase inhibitors (*44*), Aβ1-42 potentiates this marker of apoptosis. Our findings thus point to an entirely novel and selective role for the Aβ42 peptide, and raise the intriguing possibility that compromised γ-secretase activity against the CTFs of APP and/or other substrates contributes to the pathogenesis of AD.

## RESULTS

### Aβ1-42 inhibits γ-secretase-mediated proteolysis of APP_C99_

We have shown that Aβ peptides displaying a relatively low affinity for γ-secretase, when present at high concentrations, can compete with the higher affinity APP_C99_ substrate for binding to the enzyme (*30*). Based on this observation, we hypothesized that an increase in the concentration of Aβ42 may promote the establishment of an inhibitory mechanism that involves the formation of ‘non-productive’ enzyme-Aβ42 complexes. The Aβ42-mediated inhibition would result in the accumulation of unprocessed γ-secretase substrates and reduced production of their intracellular domains, both of which may contribute to dysregulated downstream signaling.

As a first step, we investigated the effects of human Aβ1-42 (from now on referred to as Aβ42) on the processing of APP_C99_ in well-controlled kinetic analyses **(Figure 1B)** (*30*). We incubated purified (wildtype) γ-secretase enzyme with purified APP_C99_-3xFLAG substrate at the K_M_ or saturating concentrations (0.4 μM and 1.2 μM, respectively) in the presence of human Aβ42 peptides at concentrations ranging from 0.5 μM to 10 μM. We then analyzed the *de novo* generation of AICD-3xFLAG by quantitative western blotting **(Supplementary figure 1A)**. Methanol:chloroform extraction was performed to remove the excess of unprocessed substrates, the high levels of which could preclude the quantitative analysis of ICDs. The analysis demonstrated that human Aβ42 inhibited γ-secretase-mediated proteolysis of APP_C99_ in a dose dependent manner. The more pronounced peptide-driven inhibition at the lower substrate concentration (IC_50_=1.3 μM vs 3.6 μM for 0.4 and 1.2 μM APP_C99_, respectively) is consistent with a mechanism in which Aβ42 competes with the substrate for the enzyme. In addition, mass spectrometry analyses of the proteolytic reactions showed that Aβ42 partially inhibited the generation of both AICD product types (AICD_50-99_ and AICD_49-99_) **(Figure 1C)**, indicating that Aβ42 inhibits both γ-secretase product lines. Next, we tested whether human Aβ42 exerted inhibition on all members of the γ-secretase family – i.e. irrespective of the type of PSEN (1 vs 2) and APH1 (A vs B) subunits **(Figure 1D)**. Quantitative western blotting analysis revealed a marked inhibition of total AICD production by all types of γ-secretases in the presence of 3 µM human Aβ42. These findings support a competitive mechanism wherein low-affinity substrates (acting also as products) are able to re-associate with the protease and inhibit the processing of transmembrane substrates when present at relatively high concentrations.

To gain further insights, we investigated γ-secretase mediated processing of Aβ42 to Aβ38 under the conditions used to examine APP_C99_, using the latter as a positive control **(Figure 1E)**. Despite the use of relatively high concentrations of Aβ42 (10 µM), this peptide was not converted into Aβ38. In contrast, proteolytic reactions using APP_C99_ (1.5 μM) resulted in the generation of Aβ38 (0.5-1 nM). We also tested whether Aβ42 served as a substrate in conditions that mimic a native-like environment, i.e. detergent resistant membranes (DRMs) **(Figure 1E)** (*30, 45*). As in the detergent conditions, Aβ42 was barely converted into Aβ38. We note that γ-secretase processes Aβ43 into Aβ40 under similar conditions, even when this peptide was added at much lower concentrations (0.5-1 µM) (*30*). Taken together, these observations indicate that exogenous Aβ42 does interact with γ-secretases but, unlike Aβ43, does not act as a substrate (at least under these conditions), supporting the notion that Aβ42-driven inhibition of γ-secretases is mediated via the formation of non-productive enzyme-substrate (E-S) like complexes. However, a scenario wherein Aβ42 interacts with APP_C99_ to reduce the amount of free APP_C99_ substrate available for the enzymatic cleavage is not excluded by these data.

We also investigated whether the inhibitory effects of Aβ42 on γ-secretase were reversible. To this end, we conjugated purified γ-secretase complexes to beads using a high-affinity anti-NCSTN nanobody and incubated the enzyme-conjugated beads with 0.4 µM APP_C99_, in the absence or presence of 3 µM Aβ42, for 40 minutes at 37°C. Note that this concentration of peptide substantially inhibited AICD generation **(Figure 1B)**. As a control, 10 µM γ-secretase inhibitor X (GSI, Inh X) was included. After the incubation, we collected the supernatants, washed the beads in assay buffer and re-incubated them with 0.4 µM APP_C99_ for 40 minutes at 37°C. Analysis of the levels of the *de novo* generated AICD products in the supernatant fractions collected before (reaction 1) and after washes (reaction 2) indicated that Aβ42 inhibition of γ-secretase is fully reversible **(Figure 1F)**. Collectively, our analyses support a model wherein Aβ42 forms a non-productive E-S like complex with γ-secretase and its binding is reversible.

### The N- and C-termini of Aβ play key roles in the inhibition of γ-secretase activity

We then investigated the structure-function relationships relevant to the Aβ42-driven inhibitory mechanism. The effects of mouse/rat (murine) Aβ42 and N-terminally truncated human Aβx-42 (11-42 and 17-42) peptides on γ-secretase activity were examined in cell-free assays using peptide concentrations ranging from 0.5 μM to 10 μM **(Figure 2A-C)**. Quantification of the *de novo* AICD product levels showed that murine Aβ42 maximally inhibited γ-secretase activity by ∼20% **(Figure 2A)**. As three amino acids in the N-terminal domain (R5G, Y10F and H13R) differentiate human and murine Aβ1-42 peptides **(Figure 1A)**, the differences in the inhibition thus defined the N-terminal domain of Aβ as contributing to the inhibitory mechanism. It is noteworthy that similar to human Aβ1-42, murine Aβ1-42 was not processed to Aβ1-38 **(Supplementary figure 1B)**. The analyses of other naturally occurring N-terminally truncated Aβx-42 peptides, generated by β-secretase (alternative) cleavage at the position 11 or by α-secretase cut at the position 17 in the Aβ sequence, showed that the truncated peptides exhibited reduced inhibitory potencies relative to Aβ42. The IC_50_ values for Aβ11-42 were reduced 1.79- and 1.31-fold (K_M_ and saturating substrate concentrations, respectively), relative to Aβ42 **(Figure 2B, Supplementary Table 1)**, while the larger N-terminal truncation (of residues 1-16) even further reduced the inhibitory effect to the level seen with murine Aβ42 **(Figure 2C)**. Collectively, these data assign a defining role to the N-terminal region of Aβ in the inhibition of γ-secretase activity.

**Figure 2.**
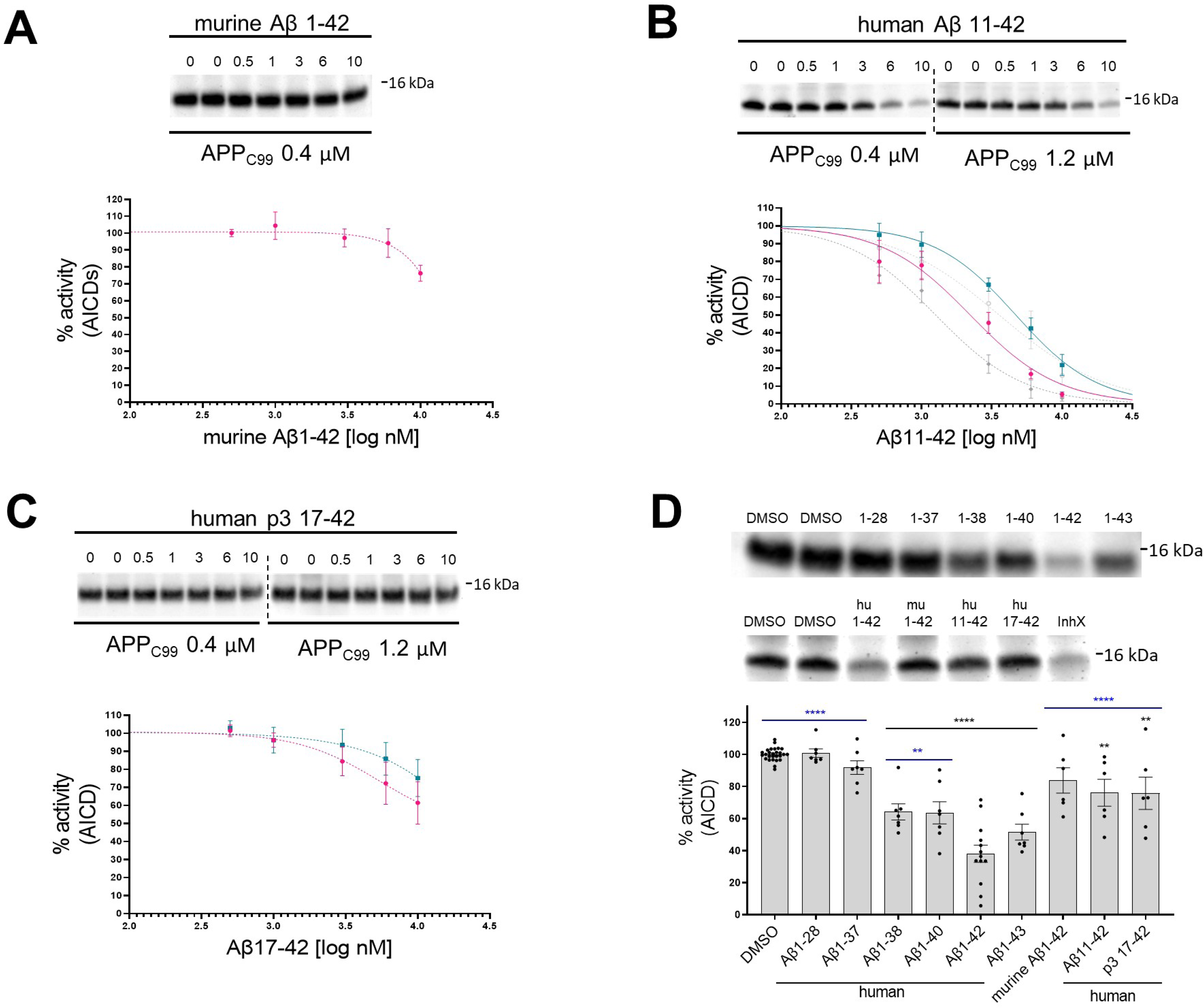
N- and C-terminus of Aβ influence the inhibitory properties of the peptide. (A, B, C) The western blots present *de novo* generated AICDs in detergent-based γ-secretase activity assays using purified protease and APP_C99_-3xFLAG at 0.4 μM or 1.2 μM as a substrate. To test the inhibitory properties of (A) murine Aβ1-42, (B) human Aβ11-42 and (C) human p3 17-42, the peptides were added to the assays at concentrations ranging from 0.5 to 10 μM. DMSO at 2.5% was used as a vehicle. The graphs present the quantification of the western blot bands for AICDs. The pink and green lines correspond to 0.4 μM and 1.2 μM substrate concentrations, respectively. The grey dotted lines on the plot B present the curves recorded for human Aβ1-42 (from Figure 1, plot B) for comparison. The data are normalized to the AICD levels generated in the DMSO conditions, considered as 100%, and presented as mean ± SEM, N=3-8. (D) Detergent-based γ-secretase activity assays using purified protease and APP_C99_-3xFLAG at 0.4 μM concentration were supplemented with different Aβ peptides at 1 μM concentration or DMSO. *De novo* generated AICDs were quantified by western blotting. The data are shown as mean ± SEM, N=6-27. The statistics were calculated using one-way ANOVA and multiple comparison Dunnett’s test, with DMSO (black) or Aβ1-42 (blue) set as references. **p<0.01, **** p<0.0001.

We next tested whether longer Aβ peptides (>Aβ42), which form more stable interactions with the protease (*30*), also inhibit γ-secretase activity. In addition, we investigated whether shortening of the C-terminus diminishes the inhibitory properties. To this end, we evaluated a series of naturally occurring Aβ1-x peptides, ranging from 37 to 43 amino acids in length, at 1 µM final concentration for their effect on γ-secretase proteolysis of APP_C99_ **(Figure 2D)**. In these experiments, the reduction in DMSO (vehicle) concentration to 0.2% was accompanied by ∼1.7 fold enhanced Aβ-mediated inhibition, relative to the conditions shown in Figure 1B. We speculate that increased Aβ potency under these conditions is explained by a modest but measurable reduction in the proteolysis by the higher concentration of DMSO used in Figure 1 assays (*46*). Relative to Aβ42, shorter Aβ species exerted progressively less or no inhibition on the *de novo* generation of AICD. It is noteworthy that peptides longer than Aβ42 serve as substrates for γ-secretases under these conditions, and their shortening potentially converts them into less potent inhibitory species (*30*). Taken together, these data strengthen the conclusion that human Aβ42 inhibits γ-secretases and indicate that both Aβ C- and N-termini modulate the inhibitory mechanism.

### Aβ42 treatment leads to the accumulation of APP C-terminal fragments in neuronal cell lines and human neurons

We reasoned that human Aβ42-driven inhibition of γ-secretase activity would lead to the accumulation of unprocessed γ-secretase substrates in cells (e.g. APP-CTFs in the case of APP) as well as reduced generation of products (Aβ peptides in the case of APP). To test these possibilities, we treated human neuroblastoma SH-SY5Y, rat pheochromocytoma PC12 and human neural progenitor cells (ReNcell VM) with recombinant human Aβ42 or p3 17-42 peptides at 1 μM or 2.5 μM final concentrations, and analyzed the levels of endogenous APP-CTFs and full-length APP (APP-FL) by western blotting **(Figure 3A-C)**. Treatment with GSI (InhX) at 2 μM was included as a positive control. Human Aβ42 treatment increased the APP-CTF/FL ratio, a read-out of γ-secretase activity. In contrast, treatment with p3 peptide had no effect. The increments in the APP-CTF/FL ratio suggested that Aβ42 (partially) inhibits the global γ-secretase activity. To further investigate this, we measured the direct products of the γ-secretase mediated proteolysis of APP. Since the detection of the endogenous Aβ products via standard ELISA methods was precluded by the presence of exogenous human Aβ42 (treatment), we used an N-terminally tagged version of APP_C99_ and quantified the amount of total secreted Aβ, which is a proxy for the global γ-secretase activity. Briefly, we overexpressed human APP_C99_ N-terminally tagged with the short 11 amino acid long HiBiT tag in human embryonic kidney (HEK) cells, treated these cultures with human Aβ42 or p3 17-42 peptides at 1 μM, or DAPT (GSI) at 10 μM, and determined total HiBiT-Aβ levels in conditioned media (CM). DAPT fully inhibits γ-secretase at the concentration used, and hence the values measured in DAPT treated conditions were used for the background subtraction. We found a ∼50% reduction in luminescence signal, directly linked to HiBiT-Aβ levels, in CM of cells treated with human Aβ42 and no effect of p3 peptide treatment, relative to the DMSO control **(Figure 3D)**. The observed reduction in the total Aβ products is consistent with the partial inhibition of γ-secretase by Aβ42.

**Figure 3.**
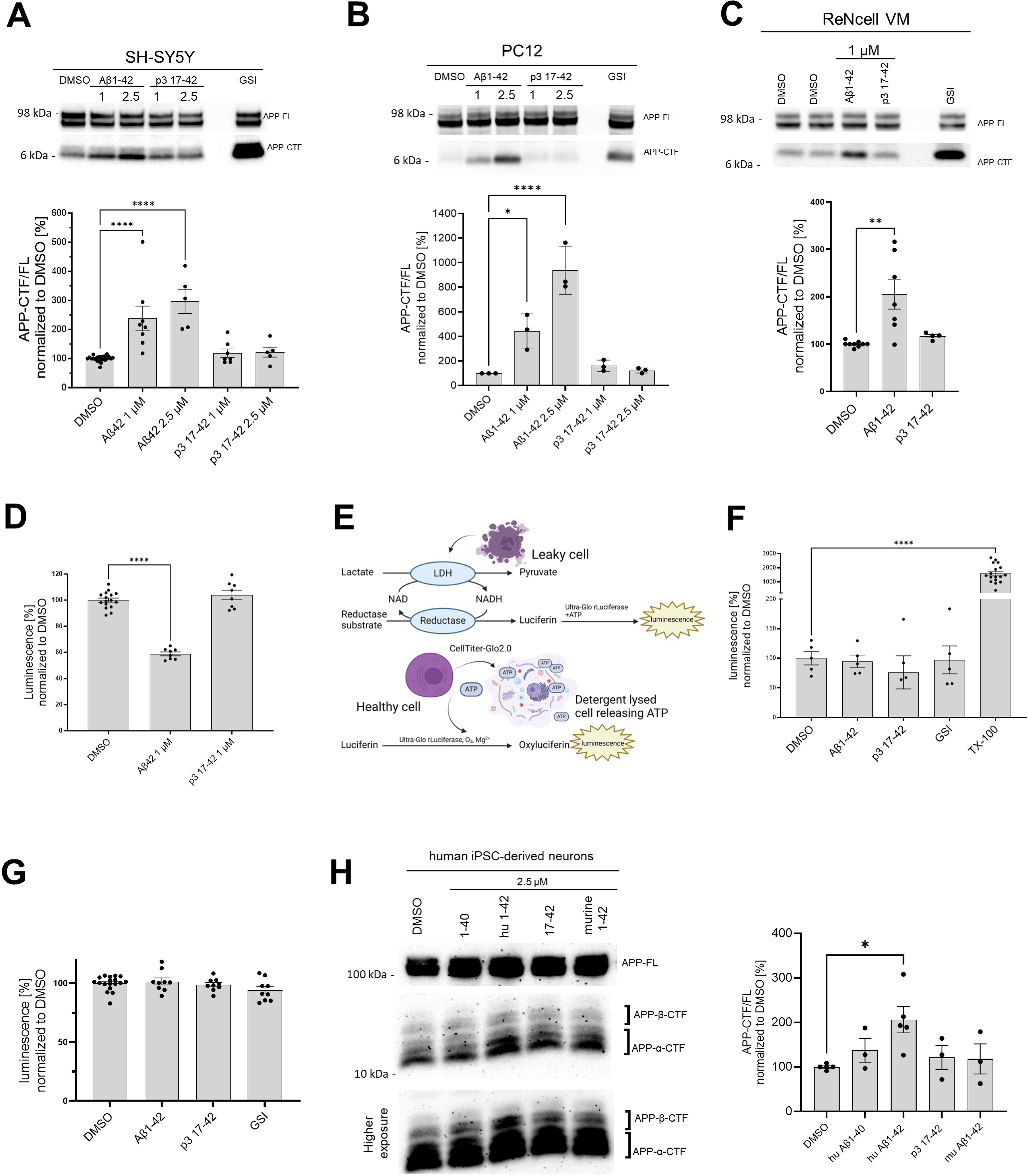
Human Aβ42 leads to the accumulation of APP-CTFs. (A, B, C, H) The western blots present full length APP (APP-FL) and APP C-terminal fragments (APP-CTFs) detected in (A) SH-SY5Y, (B) PC12, (C) ReNcell VM human neural progenitor cells and (H) induced pluripotent stem cell-derived human neurons treated for 24h with respective peptides at indicated concentrations or vehicle (DMSO). The ratio between the APP-CTF and APP-FL levels was calculated from the integrated density of the corresponding western blot bands. The data are shown as mean ± SEM, N=3-23. The statistics were calculated using one-way ANOVA and multiple comparison Dunnett’s test, with DMSO set a reference. * p<0.05, **p<0.01, **** p<0.0001. (D) The amount of HiBiT-Aβ peptides was measured in the conditioned medium collected from HEK cell line stably expressing HiBiT-APP C99 treated with DMSO, Aβ1-42 (1 μM) or p3 17-42 (1 μM). The data are shown as mean ± SEM, N=8-16. The statistics were calculated using one-way ANOVA and multiple comparison Dunnett’s test, with DMSO set a reference. **** p<0.0001. (E) The scheme presents the principles of the lactate dehydrogenase (LDH)-based and ATP-based cytotoxicity assays. The figure was created with BioRender.com. (F) Cytotoxicity of the treatments was analyzed in SH-SY5Y cells. The cells were treated with DMSO, Aβ1-42 (1 μM), p3 17-42 (1 μM) or GSI (InhX, 2 μM) for 24h, conditioned medium collected and subjected to the measurement of LDH activity using luminescence-based assay. TX-100 was used as a positive control expected to lead to 100% cell death. The data are shown as mean ± SEM, N=5-17. The statistics were calculated using one-way ANOVA and multiple comparison Dunnett’s test, with DMSO set as a reference, **** p<0.0001. TX-100 led to a marked increase in the luminescent signal, while no significant toxicity of the other treatments was detected. (G) An ATP-based cell viability assay was used to determine the cytotoxicity of the treatments. We analyzed SH-SY5Y cells treated with DMSO, Aβ1-42 (1 μM), p3 17-42 (1 μM) or GSI (Inh X, 2 μM) for 24h. The data are shown as mean ± SEM, N=9-18. The statistics were calculated using one-way ANOVA and multiple comparison Dunnett’s test, with DMSO set as a reference. No significant toxicity of the treatments was detected.

In addition, to controlling for overall cellular toxicity of Aβ42, we performed two cell toxicity assays that rely on different biological principles **(Figure 3E)**. In the first one, we quantified the activity of lactate dehydrogenase (LDH) released by cells into the conditioned medium upon plasma membrane damage **(Figure 3F)**. In the second, we detected cellular ATP as a reporter of viability. There was no significant change in these measures in response to Aβ42 (1 μM), p3 17-42 (1 μM) or GSI (2 μM) **(Figure 3G)**.

We next asked if the effects of Aβ42 would be also registered in wild type human neurons. We thus examined neurons derived from induced pluripotent stem cell (iPSC) and treated with human or murine Aβ42, p3 17-42 or human Aβ40 (all at 2.5 μM). Only treatment with human Aβ42 resulted in a significant increase of APP-CTFs **(Figure 3H)**, supporting the inhibitory role of Aβ42.

### Selective accumulation of Aβ42 in cells leads to increased levels of APP C-terminal fragments

Next, we examined the effects of a broader spectrum of Aβ peptides, presenting different N- and C-termini, on APP-CTF levels in SH-SY5Y cells. In addition, we tested murine Aβ42, which failed to inhibit γ-secretase activity in cell-free assays **(Figure 4A)**. The analysis confirmed a significant ∼2.5-fold increase in the APP-CTF/APP-FL ratio in cells treated with human Aβ42 but, intriguingly, none of the other investigated peptides significantly changed APP-CTF levels.

**Figure 4.**
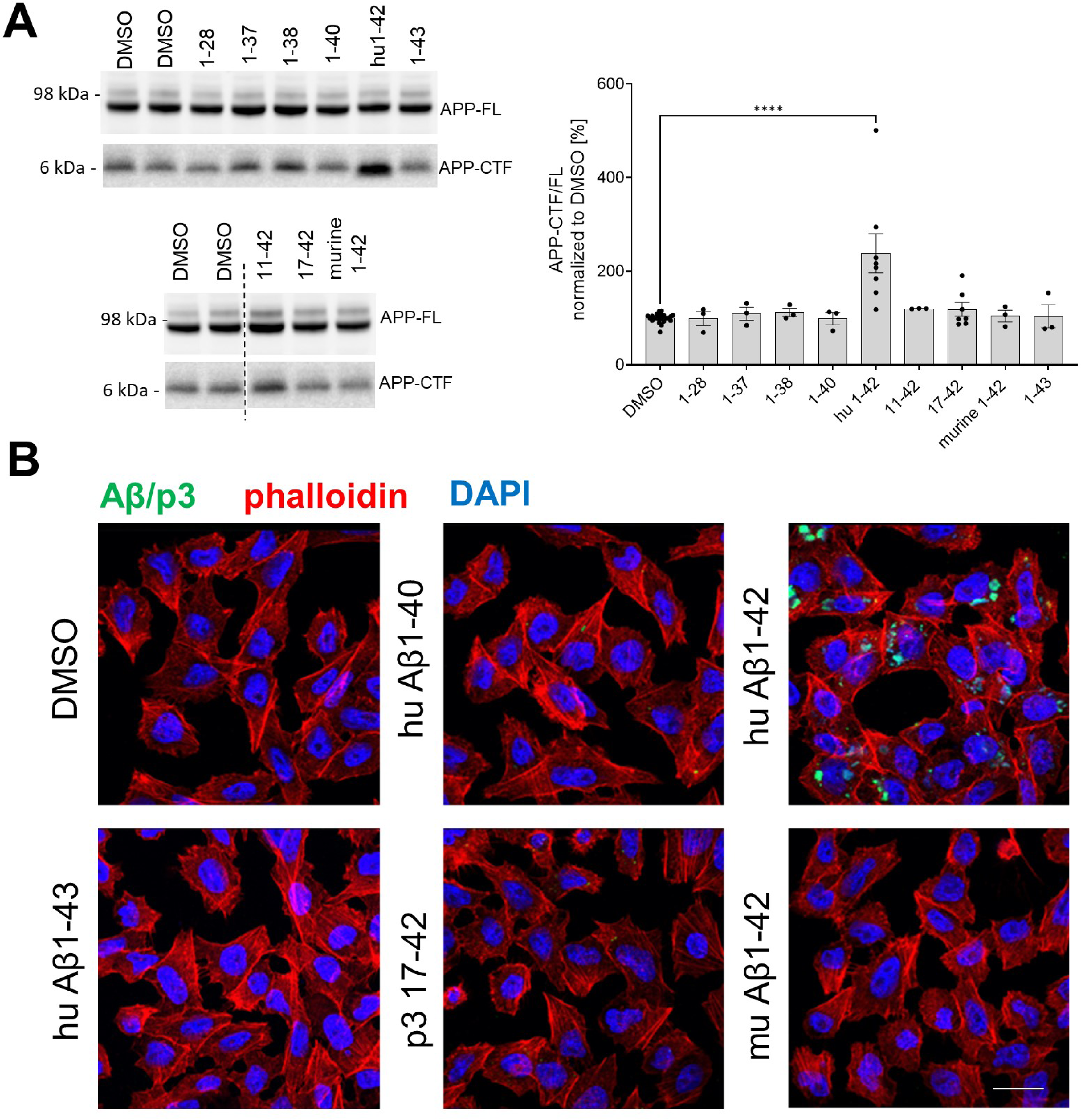
Human Aβ42 selectively accumulates in the cells and inhibits γ-secretase-mediated proteolysis. (A) APP-FL and APP-CTF levels in SH-SY5Y cells treated for 24h with a series of Aβ peptides at 1 μM concentration are shown. The APP-CTF/FL ratio was calculated from the integrated density of the corresponding western blot bands. The data are shown as mean ± SEM, N=3-23. The statistics were calculated using one-way ANOVA and multiple comparison Dunnett’s test, with DMSO set as a reference. **** p<0.0001. (B) PC12 cells were treated with respective Aβ or p3 peptides for 24h and stained with anti-Aβ antibody (clone 4G8) followed by anti-mouse Alexa Fluor Plus 488 conjugated secondary antibody, Alexa Fluor Plus 555 conjugated phalloidin and nuclear stain DAPI. Scale bar: 20 μm.

These observations provided evidence that Aβ42 differs from other tested peptides in one or more features that are critical for γ-secretase inhibition in cells. Previous studies have shown that selective cellular uptake of human Aβ42, relative to Aβ40, leading to its concentration in the acidic endolysosomal network (ELN), promotes peptide aggregation into soluble toxic Aβ species (*47–52*). We reasoned that since this compartment is the main locus of γ-secretase activity (*53*), the concentration of Aβ42 in ELN may not only promote its aggregation but also facilitate its inhibitory actions on γ-secretases. To investigate whether the selective accumulation of Aβ42 in ELN, relative to the other tested peptides, could explain the differential (cellular) inhibitory profiles **(Figure 4B)**, we treated PC12 cells with human Aβ40, Aβ42, Aβ43 or p3 17-42, or murine Aβ42 peptides at 1 µM final concentration for 24h, and examined their intracellular pools by immunostaining with an anti-Aβ/p3 antibody, 4G8 (epitope: 17-23 aa). We found that, unlike other Aβ or p3 peptides, human Aβ42 accumulated in cells. The pattern of the accumulation appeared distinct, punctate and largely perinuclear, suggestive of its presence in endolysosomal compartment. The data are consistent with a model in which intracellular accumulation of Aβ42, due to selective cellular uptake or reduced peptide degradation, distinguishes it from the other peptides studied and explains its apparently unique inhibitory properties in the cellular context.

### Human Aβ42 inhibits endogenous γ-secretase activity in neurons

We next asked whether the accumulation of APP-CTFs in cells stems from direct inhibition of γ-secretases. To answer this question, we applied an established cell-based γ-secretase activity assay to test *in situ* the protease activity in primary mouse neurons (*53, 54*) **(Figure 5A)**. This assay uses an APP_C99_-based fluorescent substrate (the C99 Y-T biosensor) to probe γ-secretase activity in living cells. This biosensor comprises APP_C99_ fused at the C-terminus with YPet, followed by a linker, Turquoise-GL and a membrane-anchoring domain to stabilize the probe in the membrane. The cleavage of the APP_C99_-based probe by endogenous γ-secretases extends the distance between YPet and Turquoise-GL, and therefore results in a decrease in the efficiency of Fӧrster resonance energy transfer (FRET) for this pair. The ratiometric nature of the reporter, and its independence of α- and β-secretase activity and cellular degradation mechanisms, allow quantitative analysis of γ-secretase activity *in situ* in living cells.

We treated mouse primary neurons with human Aβ42, p3, GSI (1 μM DAPT) or vehicle (DMSO) for 24h and performed FRET analysis **(Figure 5B-C)**. While p3 peptides did not affect the FRET signal, significant increases in FRET efficiency (i.e. increased proximity between YPet and Turquoise-GL) in cells treated with human Aβ42 or GSI, relative to vehicle treated cultures, indicated increments in the levels of the full-length (uncleaved) fluorescent substrate, and thus reduced γ-secretase activity. These results provided quantitative evidence in a cellular context for the inhibition of endogenous γ-secretases by human Aβ42, but not p3 17-42 peptides.

**Figure 5.**
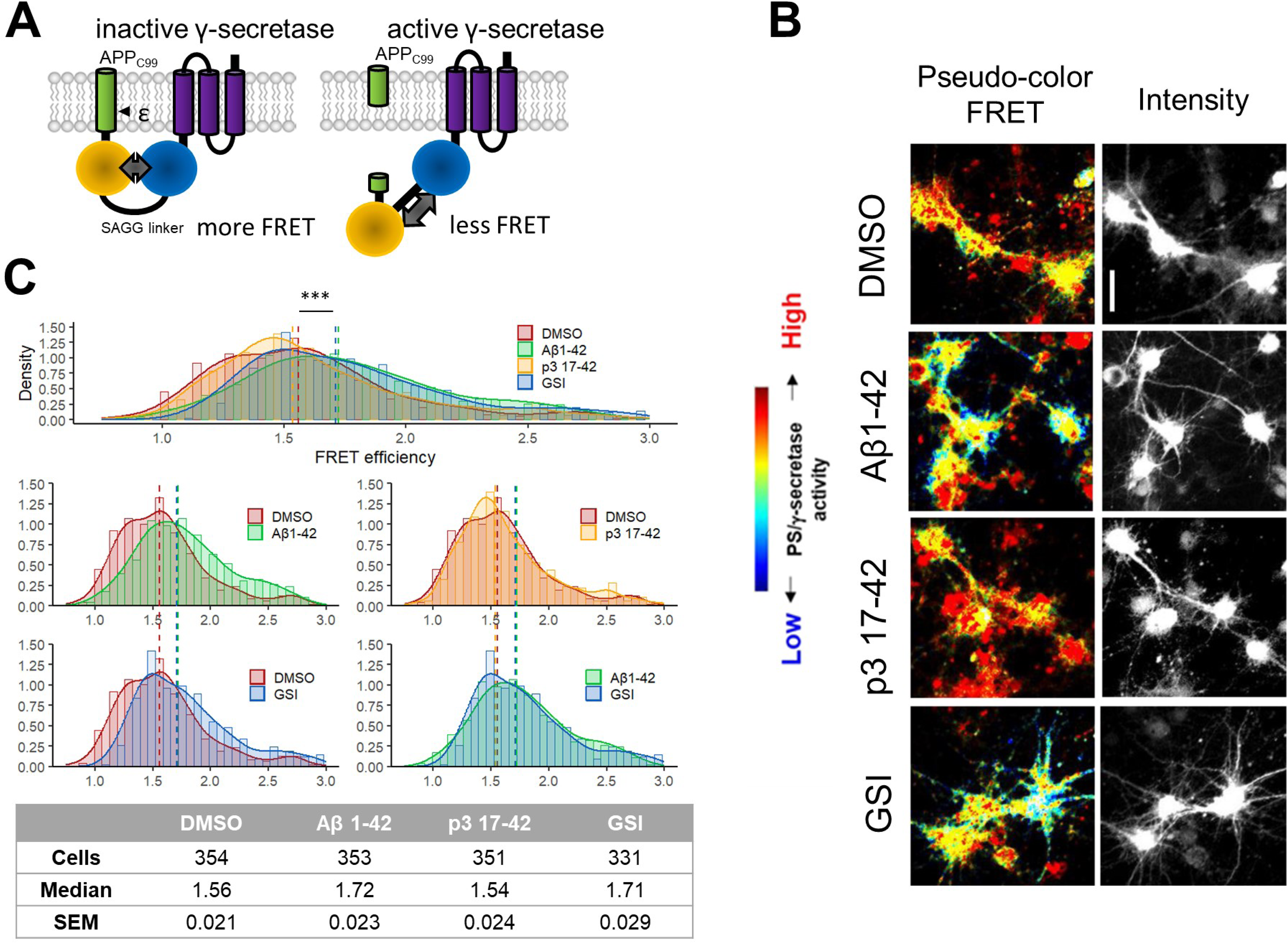
Human Aβ42 inhibits endogenous γ-secretase activity in neurons. (A) The scheme presents the FRET-based probe allowing monitoring of γ-secretase activity *in situ* in living cells. (B, C) Spectral FRET analysis of γ-secretase activity in mouse primary neurons using C99 Y-T probe is shown. The cells were treated with the indicated peptides/compounds at 1 μM concentration for 24h. Γ-secretase-mediated proteolysis results in an increase in the distance between two fluorophores incorporated in the probe (YPet and Turquoise-GL). The increase in the distance translates to the reduced FRET efficiency, quantified by the YPet/Turquoise-GL fluorescence ratio. The distribution of recorded FRET efficiency, inversely correlating with γ-secretase activity, is shown in the density plots, N=4 (n=331-354). Medians are shown as dashed lines. Optimal bin number was determined using Freedman-Diaconis rule. The statistics were calculated using Kruskal-Wallis test and multiple comparison Dunn test. Significant differences (*** p<0.001) were recorded for DMSO vs Aβ1-42, p3 17-42 vs Aβ1-42, DMSO vs GSI and p3 17-42 vs GSI.

### Aβ42-mediated γ-secretase inhibition is the major mechanism contributing to APP-CTF accumulation in cells

Our data support a direct effect of Aβ42 peptides on the activity of γ-secretase in well controlled, cell-free systems and in cell-based assays. Although the cell-free and FRET-based systems are independent of α- and β-secretase activity and cellular degradation pathways, we elected to further pursue the possibility that changes in the activity of these enzymes or degradation of CTFs could contribute to the observed increase in APP-CTFs in cellular systems. We investigated the impact of Aβ42 on α-secretase (ADAM10) and β-secretase (BACE1) levels, alterations in which could result in increased APP-CTF levels, by analyzing the expression of ADAM10 and BACE1 in total cell lysates from SH-SY5Y cells treated with Aβ40 (1 μM), Aβ42 (1 μM), p3 17-42 (1 μM), GSI (InhX, 2 μM) or vehicle DMSO control **(Supplementary figure 2A-B)**. We found no evidence of increased either α-secretase and β-secretase levels. In addition, we explored possible differences in the activities of these secretases by examining total soluble APP (sAPP) in conditioned medium collected from SH-SY5Y cells treated under the same conditions. Our analysis showed a reduction in total sAPP levels in the conditioned medium collected from cells treated with Aβ42. This finding is inconsistent with an Aβ42-mediated effect that would increase APP-CTF levels as result of activating α-secretase or β-secretase activity. Further studies are required to define whether the decrease in total sAPP is a consequence of sheddase inhibition or relates to an effect of Aβ42 on the rate of recycling and release of sAPP fragments. We also evaluated potential γ-secretase independent changes in the degradation of APP-CTF by assessing the impact of Aβ42 peptides on γ-secretase independent APP-CTF half-life using the well-established cycloheximide (CHX)-based assay. CHX inhibits protein synthesis by blocking translation elongation (*55*). In these experiments, we pre-treated SH-SY5Y cells with either GSI (DAPT, 10 μM) or GSI + Aβ42 (1 μM) for 24h. Importantly, the GSI treatment allowed for complete γ-secretase inhibition in both conditions, resulting in the maximum accumulation of APP-CTFs that can possibly be achieved by blocking γ-secretase processing. We then added CHX (50 μg/ml) while maintaining GSI and vehicle versus GSI plus Aβ42. In addition, we treated with bafilomycin A1 (200 nM), which de-acidifies lysosomes and compromises degradation pathways, as a positive control **(Figure 6A)**. We collected cells thereafter at 0h, 1h, 2.5h and 5h.

**Figure 6.**
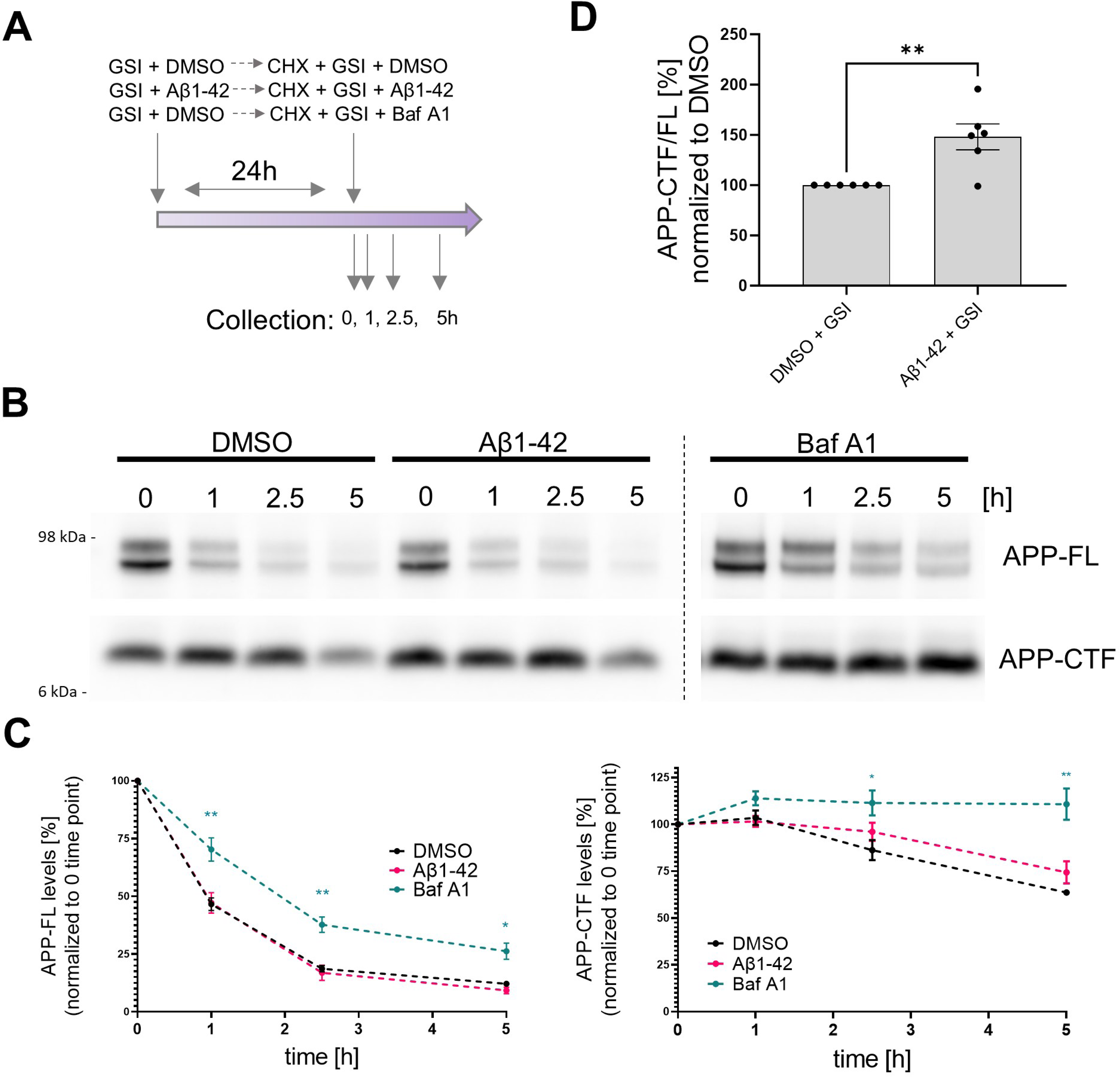
Contribution of Aβ42-mediated inhibition of γ-secretase and general degradation mechanisms to APP-CTF accumulation. (A) The scheme presents the experimental design of the cycloheximide (CHX)-based assay evaluating APP-FL and APP-CTF stability. (B) Western blot shows APP-FL and APP-CTF levels in SH-SY5Y cells at 0, 1, 2.5 and 5h collection points defined in the scheme (A). (C) The integrated densities of the bands corresponding to APP-FL and APP-CTF were quantified and plotted relatively to the time point zero. The data are presented as mean ± SEM, N=6. Statistics were calculated using two-way ANOVA, followed by multiple comparison Dunnett’s test. *p<0.05, **p<0.01. (D) Quantification of APP-CTF/ FL ratio at the zero time point is shown. The data are presented as mean ± SEM, N=6. Statistics were calculated using unpaired Student’s t-test, **p<0.01.

Quantitative analysis of the levels of both APP-CTFs and APP-FL over the 5h time-course showed that bafilomycin A1 treatment markedly prolonged the half-life of both proteins but did not reveal significant differences in their levels between GSI versus GSI plus Aβ42-treated cells **(Figure 6 B-C)**. The lack of a significant effects of Aβ42 on the levels of APP-CTFs under these conditions (full γ-secretase inhibition) suggests that the peptide does not induce major defects in cellular degradation pathways, at least at this concentration. However, we noted an increased APP-CTF/FL ratio in GSI + Aβ42 treated cells vs GSI + DMSO treated ones at time zero **(Figure 6D),** suggesting that the Aβ42-induced increase in APP-CTFs is mediated by both inhibition of γ-secretase and one or more additional mechanisms. Nevertheless, the greater relative accumulation of APP-CTFs in the absence (138%) **(Figure 3A)** versus the presence (48%) of the GSI **(Figure 6D)** is evidence that Aβ42-mediated γ-secretase inhibition plays a more prominent role.

### Aβ peptides derived from biological sources trigger APP C-terminal fragment accumulation

We next asked whether Aβ from disease-relevant biological sources would recapitulate the effects of recombinant Aβ42. For these studies we used culture medium conditioned by human iPSC-derived neurons expressing APP wild type or the KM670/671NL (SWE) variant. The APP SWE mutation increases total Aβ production by promoting the amyloidogenic processing of APP. ELISA-based quantification of Aβ content estimated a total concentration in the low nanomolar range (0.5-1 nM). Conditioned medium (WT or SWE) was applied to PC12 cells either directly or after Aβ depletion (delSWE) using the anti-APP 3D6 (epitope Aβ 1-5) **(Figure 7)**. After a 24h incubation at 37°C, the APP-CTF/FL ratio was measured. Cells incubated in control, unconditioned media were used as a reference control. We observed significant increments in the APP-CTF/FL ratio in cells treated with SWE conditioned media, relative to control cells. Aβ immunodepletion from the SWE medium (delSWE-CM) lowered the APP-CTF/FL ratio to the levels observed in the control cells. These data provide evidence that Aβ peptides derived from biological sources induce the accumulation of APP-CTFs even when present at low nM concentrations, and thus point to this biological source of Aβ as being far more potent than recombinant Aβ42.

**Figure 7.**
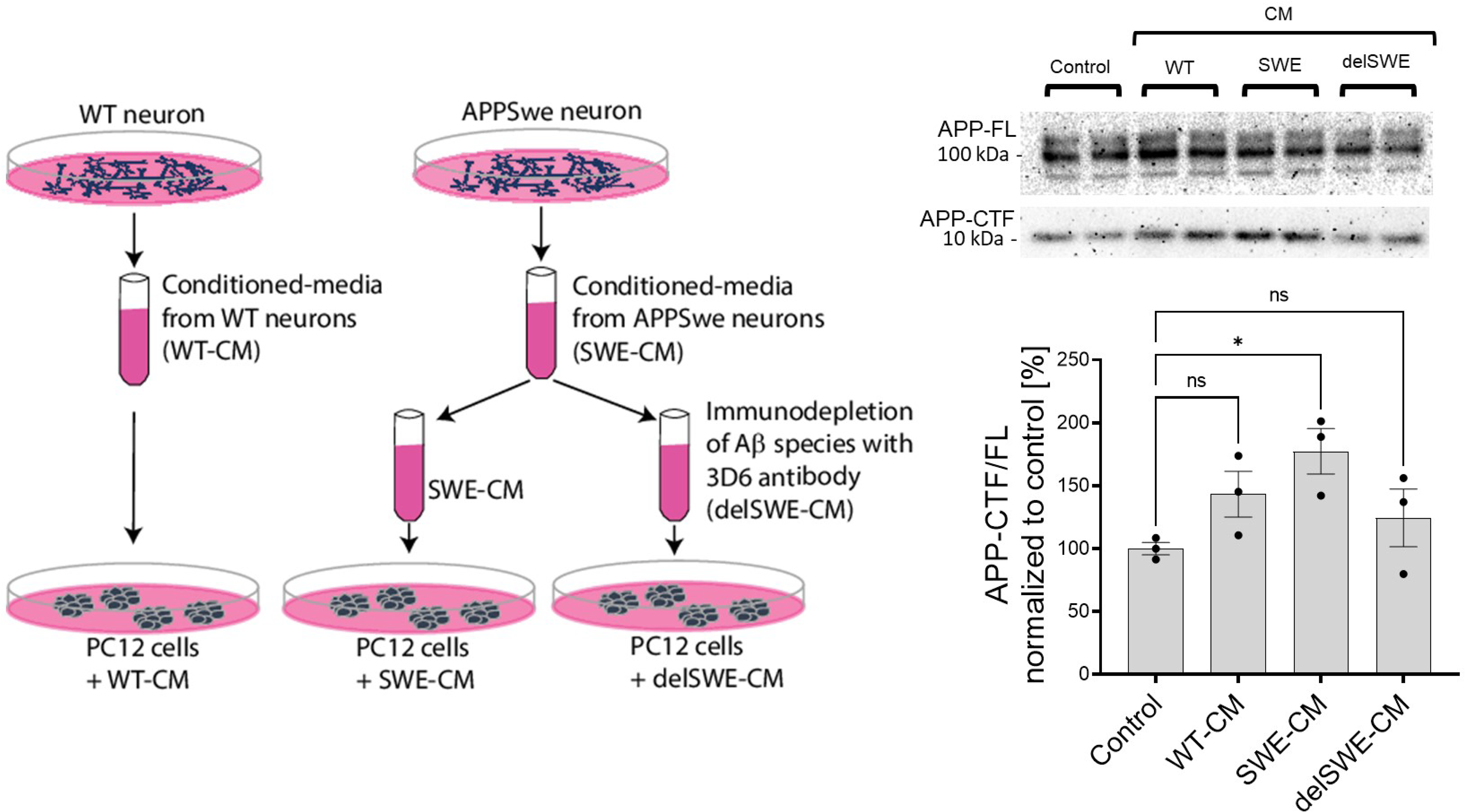
Human Aβ42 leads to the accumulation of APP-CTFs. (A) Scheme depicts the experimental design testing the impact of biologically-derived Aβ on the proteolysis of APP. Conditioned media were collected from fully differentiated WT human neurons (WT-CM) and human neurons expressing APP with Swedish mutation (SWE-CM). A portion of SWE-CM was subjected to Aβ immunodepletion using the anti-Aβ antibody 3D6. WT, SWE and Aβ immunodepleted (delSWE-CM) CMs were added onto the PC12 cells for 24h to analyze APP processing. As a reference, control cells treated with base media were analyzed. Representative western blots present the analysis of total cellular proteins from four cell culture sets treated with base media (control), WT-CM, SWE-CM and delSWE-CM, respectively. The ratio between APP-CTFs and APP-FL was calculated from the integrated density of the corresponding western blot bands. The data are shown as mean ± SEM, N=3. The statistics were calculated using one-way ANOVA and multiple comparison Dunnett’s test, with control set as a reference, *p<0.05.

### Human Aβ42 peptides inhibit proteolysis of other γ-secretase substrates, beyond APP-CTFs

The Aβ42-mediated inhibition of endogenous γ-secretase activity in cells raises the possibility that Aβ42 would inhibit the processing of γ-secretase substrates beyond APP-CTFs. To address this, we analyzed the effects of human Aβ42 on the processing of a purified NOTCH1-based substrate in cell-free conditions **(Figure 8A)**. We incubated purified γ-secretase and the NOTCH1-based substrate (at 0.4 μM or 1.2 μM concentrations) in the presence of increasing amounts of human Aβ1-42 (ranging from 0.5 μM to 10 μM). Quantification of the *de novo* generated NICD-3xFLAG showed a dose dependent inhibition of γ-secretase proteolysis of NOTCH1 by Aβ42. As for APP_C99_, the derived IC_50_ values showed that the degree of the inhibition depended on substrate concentration, consistent with a competitive mechanism **(Supplementary Table 1)**. We extended this analysis by assessing the γ-secretase-mediated proteolysis of other substrates (ERBB4-, neurexin- and p75-based) in the presence of 3 μM human Aβ42 **(Figure 8B)**. Quantification of the respective *de novo* generated ICDs revealed that Aβ42 reduced γ-secretase proteolysis of each of these substrates. Mass spectrometry-based analyses of the respective ICDs confirmed the inhibition **(Supplementary figure 3)**.

**Figure 8.**
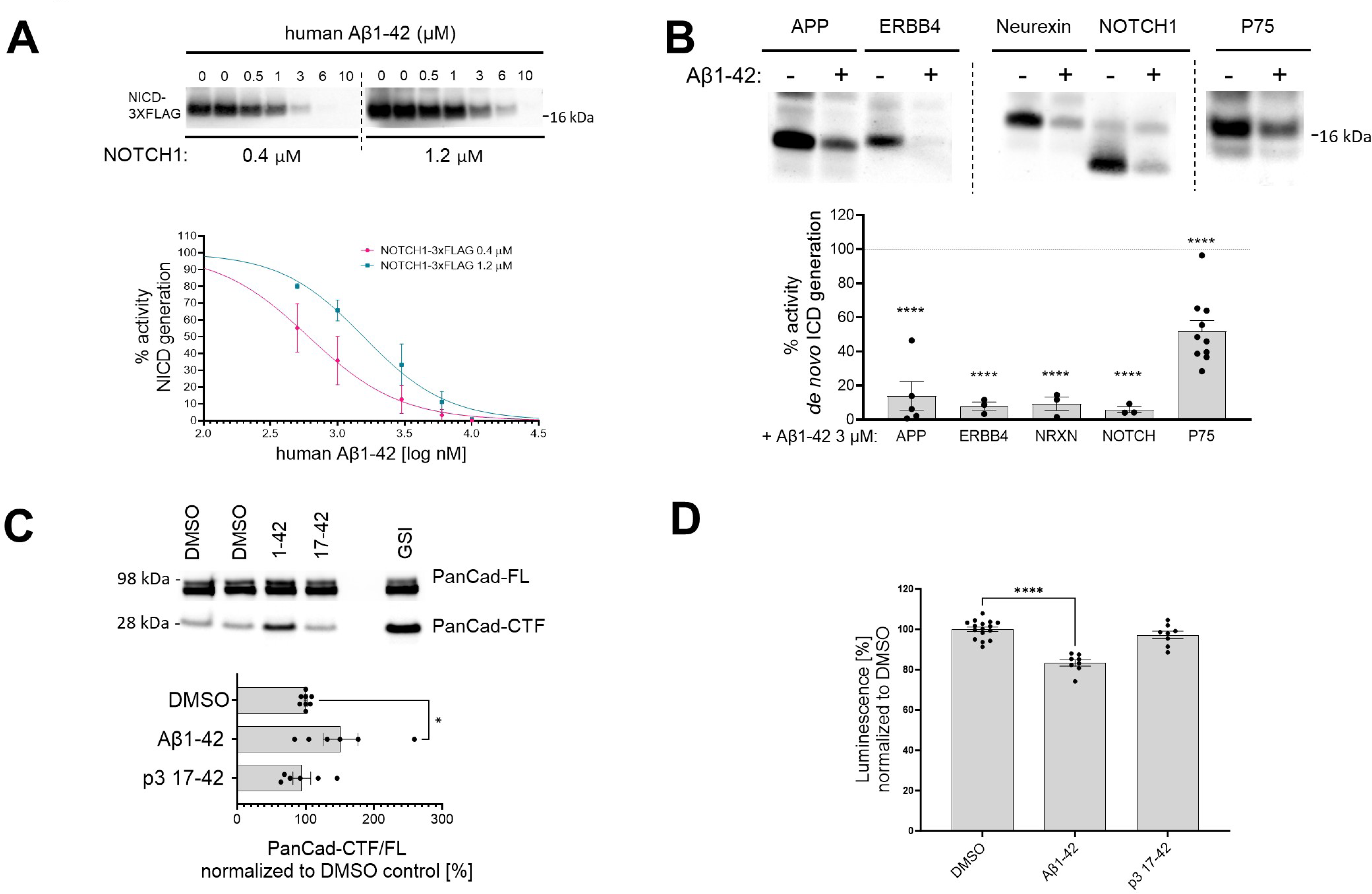
Human Aβ1-42 peptides inhibit proteolysis of multiple γ-secretase substrates. (A) The western blot presents *de novo* generated NICDs in detergent-based γ-secretase activity assays, using NOTCH1-3xFLAG at 0.4 μM and 1.2 μM as a substrate, supplemented with human Aβ1-42 peptides at concentrations ranging from 0.5 to 10 μM. The graphs present the quantification of the western blot bands for NICDs. The pink and green lines correspond to 0.4 μM and 1.2 μM substrate concentrations, respectively. The data are normalized to the NICD levels generated in the DMSO conditions, considered as 100% and presented as mean ± SEM, N=3-5. (B) Analysis of the *de novo* ICD generation in cell-free detergent-based γ-secretase activity assays is shown. The graph presents the quantification of the western blots. The data are shown as mean ± SEM, N=3-18. The statistics were calculated using one-way ANOVA and multiple comparison of predefined columns, with Šidák correction test, with respective DMSO supplemented reactions set as a reference, **** p<0.0001. (C) PanCad-FL and PanCad-CTF levels in ReNcell VM cells treated for 24h with human Aβ1-42 peptides at 1 μM or GSI (InhX) at 2 μM concentration were quantified by western blotting. The PanCad-CTF/FL ratio was calculated from the integrated density of the corresponding bands. The data are presented as mean ± SEM, N=6-8. The statistics were calculated using one-way ANOVA and multiple comparison Dunnett’s test, with DMSO set as a reference. *p<0.05. (D) The graph presents the quantification of the HiBiT-Aβ like peptide levels in conditioned media collected from HEK cell line stably expressing the HiBiT-NOTCH1 based substrate and treated with DMSO, Aβ1-42 (1 μM) or p3 17-42 (1 μM). The data are shown as mean ± SEM, N=8-16. The statistics were calculated using one-way ANOVA and multiple comparison Dunnett’s test, with DMSO set a reference. **** p<0.0001.

We next investigated the effects of human Aβ42 on γ-secretase-mediated processing of pan-cadherin in ReNcell VM cells. Cleavage of cadherins by α-secretase generates a membrane bound CTF, which is a direct substrate of γ-secretases (*56*). As before, we used the PanCad-CTF/FL ratio to estimate the efficiency of the γ-secretase-mediated proteolytic processing **(Figure 8C)**. Treatment with human Aβ42 and InhX, but not p3, resulted in significant increases in the PanCad-CTF/FL ratio, demonstrating that Aβ42 inhibits processing of several different γ-secretase substrates. These results suggest that this peptide can inhibit the proteolysis of γ-secretases substrates in general.

Finally, we measured the direct N-terminal products generated by γ-secretase proteolysis from a HiBiT-tagged NOTCH1-based substrate, an estimate of the global γ-secretase activity. We quantified the Aβ-like peptides secreted by HEK 293 cells stably expressing this HiBiT-tagged substrate upon treatment with 1 µM Aβ1-42 or p3 17-42 peptide, and 10 µM DAPT (GSI) as control for background subtraction **(Figure 8D)**. A ∼20% significant reduction in the amount of secreted N-terminal HiBiT-tagged peptides derived from the NOTCH1-based substrates in cells treated with Aβ1-42 supports the inhibitory action of Aβ1-42 on γ-secretase mediated proteolysis.

By demonstrating that Aβ42 inhibits γ-secretase proteolysis of substrates that are structurally distinct from APP-CTFs, these data indicate that Aβ42 inhibits enzyme activity independent of substrate structure. This renders much less likely the mechanistic possibility that inhibition is due to an interaction with the substrate.

### Human Aβ42 driven accumulation of p75-CTF induces markers of neuronal death

The p75 neurotrophin receptor plays a prominent role in neurotrophin signaling. Under certain conditions its C-terminal fragment (p75-CTF, a γ-secretase substrate) modulates cell death (*57, 58*). Previous studies have shown that accumulation of p75-CTF in basal forebrain cholinergic neurons (BFCNs), due to pharmacological inhibition of γ-secretases, triggers apoptosis in a TrkA activity dependent manner (*44*). The proapoptotic action of p75-CTF is prevented by expression of the TrkA receptor for nerve growth factor (NGF), which serves as a co-receptor with p75 (*44*). Similar observations have been made in PC12 cells, which model sympathetic neurons. In this case, γ-secretase inhibitors increased apoptosis in TrkA-deficient (PC12nnr5) (*59*), but not in wild type, TrkA expressing cells. These findings provided an experimental system to investigate the effects of Aβ42-mediated inhibition on γ-secretase-mediated signaling in neurons and neuron-like cells.

We tested the impact of Aβ on p75 signaling in wild type and PC12nnr5 cells. We exposed the cells to 1 µM human Aβ42, 1 µM p3, GSI (10 µM compound E) or vehicle (DMSO) for 3 days, and measured apoptosis via immunofluorescence staining for cleaved caspase 3 **(Figure 9A)**. In addition, to assess γ-secretase proteolysis of p75, we measured the p75-CTF/FL ratio in the treated cultures **(Figure 9B)**. Cleaved caspase 3 staining was increased in PC12nnr5 cells treated with GSI or human Aβ42, but not in those exposed to p3 or vehicle (DMSO). Of note, and consistent with our previous findings (*44*), none of the treatments elicited significant increases in cleaved caspase 3 in wild type PC12 cells that express physiological levels of TrkA. In addition, analysis of the p75-CTF/FL ratio demonstrated higher p75-CTF levels in GSI and Aβ42 treated cells. These results, together with our previous observations, strongly suggest that Aβ42-mediated accumulation of p75-CTFs results in cell death signaling in the absence or near absence of TrkA expression. Our similar studies using BFCNs recapitulated the reported increased cell death under conditions in which both γ-secretase and TrkA activities were inhibited (CE and K252α inhibitor) (*44*), and showed that treatment with 1 µM human Aβ42 mimics the increase in cleaved caspase 3 driven by the GSI **(Figure 9C)**. In conclusion, these results are evidence of a novel role for human Aβ42 peptide in the global inhibition of γ-secretase activity and dysregulation of cellular homeostasis.

**Figure 9.**
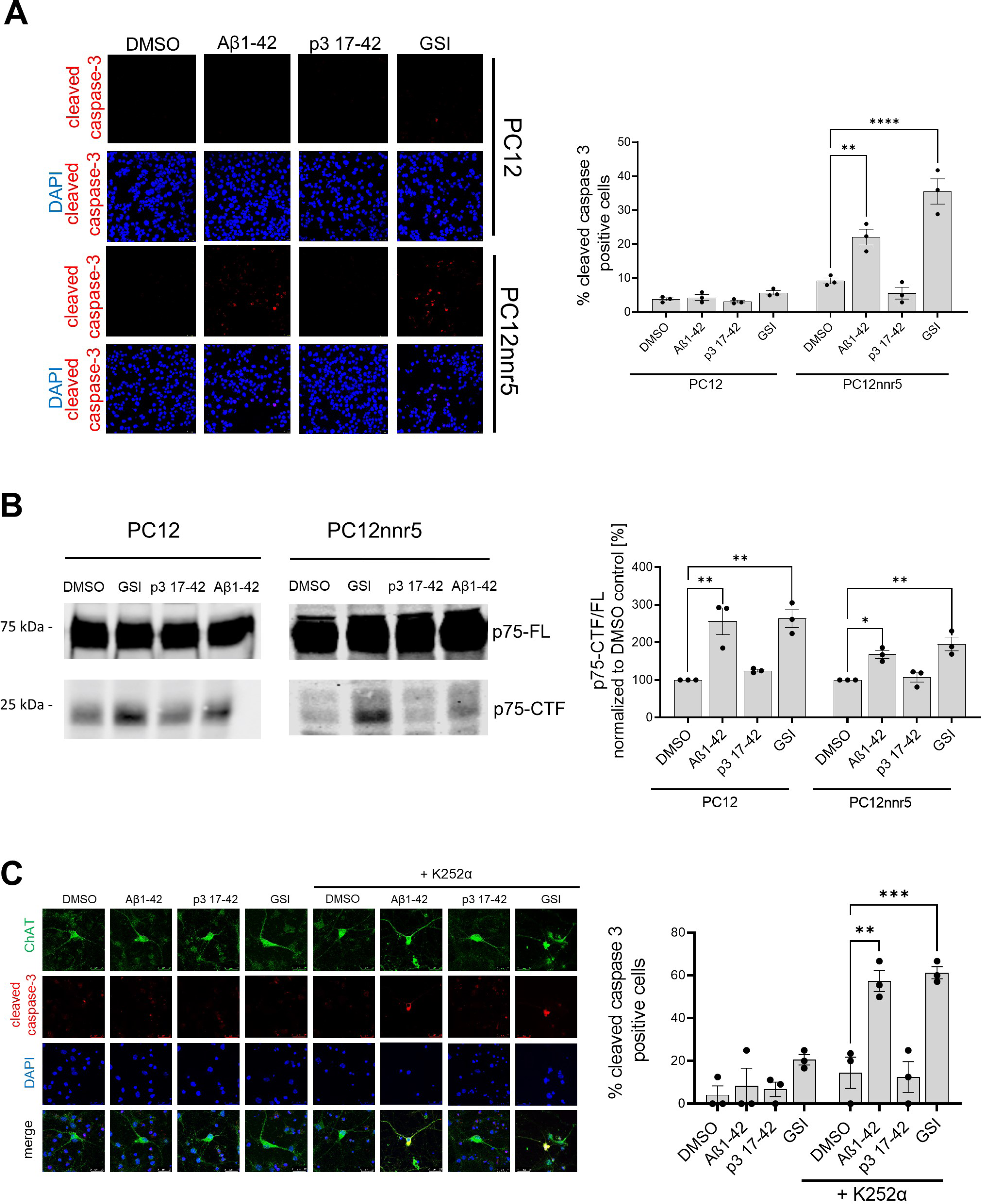
Human Aβ1-42 peptides compromises p75-mediated signaling. (A) PC12 wild type or PC12 deficient for TrkA (PC12nnr5) were incubated with human Aβ1-42 or p3 17-42 peptides, GSI (compound E) or vehicle (DMSO) for 72h. The images present immunocytochemical analyses of cleaved caspase 3, and the graph corresponding quantification of the percentage of cleaved caspase 3 positive cells. The statistics were calculated within PC12 and PC12nnr5 groups using one-way ANOVA and multiple comparison Dunnett’s test, with DMSO set as a reference. The data are presented as mean ± SEM, N=3. **p<0.01,****p<0.0001. (B) Representative western blot demonstrates the accumulation of p75-CTFs in the cells treated with human Aβ1-42 peptide or GSI. The data are presented as mean ± SEM, N=3. The statistics were calculated within PC12 and PC12nnr5 groups using one-way ANOVA and multiple comparison Dunnett’s test, with DMSO set as a reference. *p<0.05, **p<0.01. (C) Mouse primary neurons were treated with human Aβ1-42 (1 μM), p3 17-42 (1 μM), GSI (compound E, 10 μM) or vehicle control, in the absence or presence of K252α inhibitor at 0.5 μM. Level of apoptosis in basal frontal cholinergic neurons (BFCNs) was analyzed by immunostaining for choline acetyltransferase (ChAT) and cleaved caspase 3. Representative images are shown. The graph presents the quantification of the percentage of cleaved caspase-3 positive cells among ChAT positive cells. The data are presented as mean ± SEM, N=3. The statistics were calculated within -K252α and +K252α groups using one-way ANOVA and multiple comparison Dunnett’s test, with DMSO set as a reference. **p<0.01, ***p<0.001.

### APP-CTF levels are increased in Aβ42-treated synaptosomes derived from mouse brains

Since both the amyloidogenic processing of APP and the accumulation of Aβ occur at the synapse (*60, 61*), we reasoned that synaptosomes are potentially a cellular locus where elevated levels of Aβ could be acting on γ-secretase. We thus tested whether γ-secretase-mediated processing of APP at the synapse is inhibited by Aβ42 **(Figure 10A)**. To this end, we prepared synaptosomes from wildtype mouse brains, treated these fractions with 2.5 µM Aβ42 or vehicle, and then measured APP-FL and APP-CTF levels by western blotting. An increased APP-CTF/FL ratio in Aβ42 treated samples, compared to vehicle control, provided evidence of γ-secretase inhibition.

**Figure 10.**
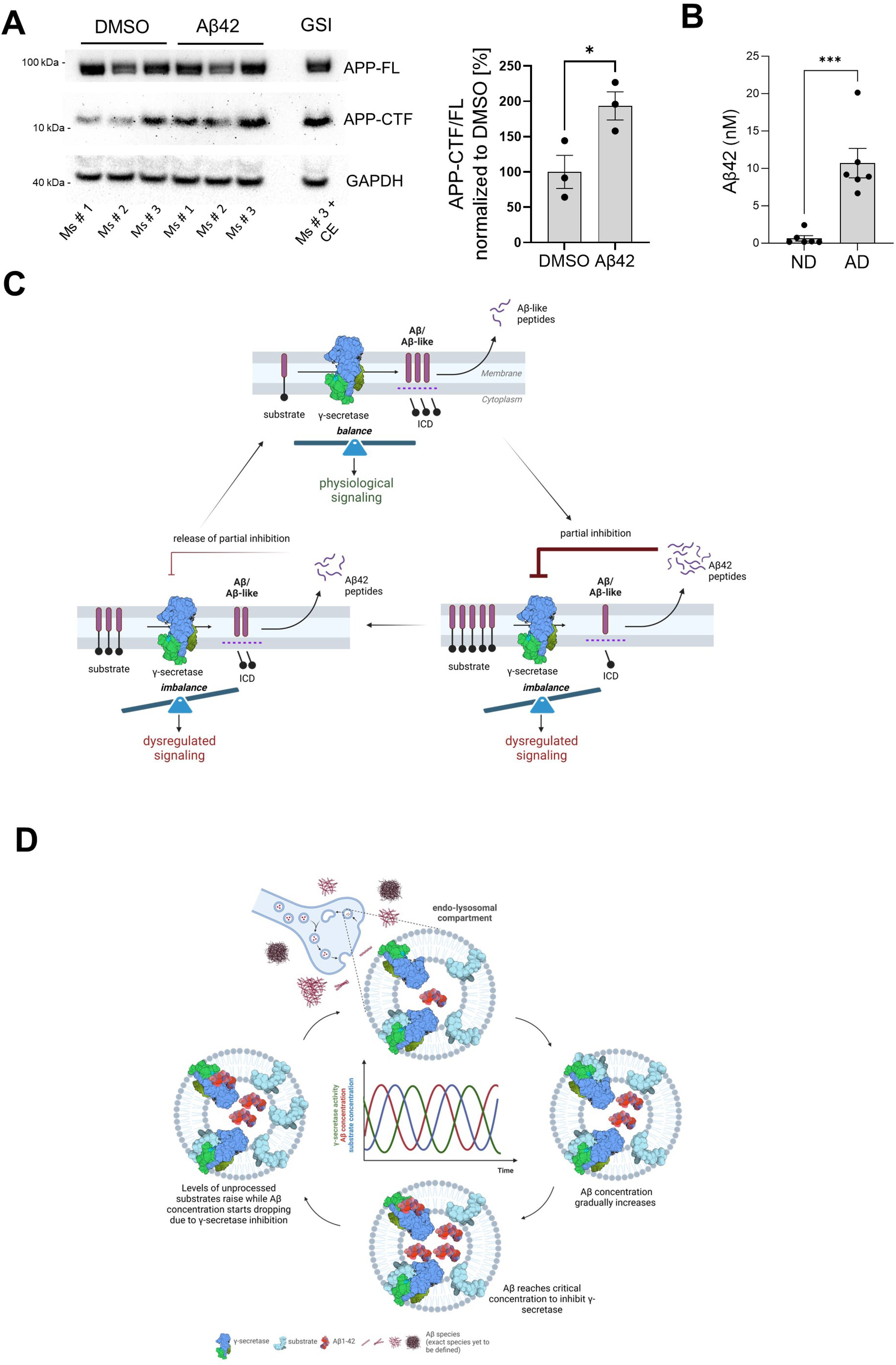
Aβ42 accumulates in AD-linked synaptosomes and treatment with this peptide increases APP-CTFs in control, mouse-derived synaptosomes. (A) Synaptosomes from brain cortices of three wild type mice were isolated, treated with DMSO or Aβ1-42, and analyzed by western blotting for APP-FL and APP-CTFs. Note the increase in APP-CTF/ FL ratio in Aβ1-42 treated samples relative to the control. The data are presented as mean ± SEM, N=3. The statistics were calculated using unpaired Student’s t-test. *p<0.05. (B) Aβ42 levels in synaptosomes derived from frontal cortices of post-mortem AD and age-matched non-demented (ND) control individuals were measured by ELISA. The data are presented as mean ± SEM, N=3. The statistics were calculated using unpaired Student’s t-test. *** p<0.001. (C) Schematic representation of the Aβ-driven γ-secretase inhibitory feedback model is shown. Figure was created with BioRender.com. (D) Scheme of the inhibitory model. Pathological increments in Aβ42 and endosomal accumulation of the peptide facilitate the establishment of an inhibitory product feedback mechanism that results in impairments in γ-secretase-mediated homeostatic signaling and contributes to AD progression. The inhibitory mechanism is complex, competitive and reversible, and hence results in pulses of γ-secretase inhibition. Figure was created with BioRender.com.

We next investigated the levels of Aβ42 in synaptosomes derived from frontal cortices of post-mortem AD and age-matched non-demented (ND) control individuals **(Figure 10B)**. Towards this, we prepared synaptosomes from frozen brain tissues using Percoll gradient procedure (*62, 63*). Intact synaptosomes were spun to obtain a pellet which was resuspended in minimum amount of PBS, allowing us to estimate the volume containing the resuspended synaptosome sample. This is likely an overestimate of the actual synaptosome volume. Finally, synaptosomes were lysed in RIPA buffer and Aβ peptide concentrations measured using ELISA (MSD). We observed that the concentration of Aβ42 in the synaptosomes from (end-stage) AD tissues was significantly higher (10.7 nM) than those isolated from non-demented tissues (0.7 nM), p<0.0005***. These data provide evidence for accumulation at nM concentrations of endogenous Aβ42 in synaptosomes in end-stage AD brains. Given that we measured Aβ42 concentration in synaptosomes, we speculate that even higher concentrations of this peptide may be present in the endolysosome vesicle system, and therein inhibit the endogenous processing of APP-CTF at the synapse. Of note treatment of PC12 cells with conditioned medium containing even lower amounts of Aβ (low nanomolar range (0.5-1 nM)) resulted in the accumulation of APP-CTFs.

## DISCUSSION

Compelling evidence supports that Aβ peptides trigger molecular and cellular cascades leading to neurodegeneration (*64, 65*). Here, we discovered and extensively characterized a previously unreported role for Aβ: inhibition of the activity of γ-secretase complexes **(Figure 10C and 10D)**. The recognition that Aβ peptides, despite lower affinities, can compete with APP-CTFs for binding to γ-secretase, when present at relatively high concentrations (*30*), led us to propose a mechanism that connects increases in the Aβ42 peptide with the inhibition of γ-secretases and dysregulation of signaling cascades relevant to neuronal physiology. In this regard, we note that the inhibition of γ-secretase in the adult mouse brain leads to age-dependent neurodegenerative phenotypes (*8–11*) by poorly understood but APP-independent (*8*) mechanisms, whereas pharmacological inhibition of γ-secretase has been linked to cognitive worsening in AD affected individuals (*7*). Here, we found *in vitro* that human (not murine) Aβ42 can inhibit γ-secretase processing, resulting in *i)* the accumulation of substrates at the membrane, *ii)* reductions in the release of soluble intracellular fragments from substrates, and *iii)* dysregulation of γ-secretase dependent signaling.

Our analysis of the global γ-secretase (endopeptidase) activity in well-controlled cell-free assays, using purified protease and substrate, demonstrated that human Aβ42 inhibited the processing of APP_C99_ by all members of the human γ-secretase family. Notably, despite the high homology between human and murine Aβ sequences, murine Aβ1-42 failed to inhibit the proteolysis of APP_C99_, implying that structural determinants in the N-terminal domain of Aβ are involved in the inhibitory mechanism. Consistent with this view, N-terminally truncated Aβ peptides (Aβ11-42 and p3 17-42) demonstrated reduced or no inhibitory properties, compared to human Aβ42. The p3 peptides lack the 16 amino acid long, hydrophilic and disordered N-terminal domain present in Aβ, but do contain the two aggregation prone regions (16–21 and 29-42) required for the assembly of oligomers (*66, 67*) and fibrils (*68–70*). Despite p3 peptides’ aggregation prone behavior (*22*) and their accumulation in AD brain (*68, 71–73*), the neurotoxicity of these peptides is not well defined (*37*). Moreover, our analyses in cell-free systems revealed that the C-terminus of Aβ also modulates its intrinsic inhibitory properties, with shorter Aβ1-x species (x= 37, 38, 40) either not acting as inhibitors or inhibiting the protease to a lesser degree than human Aβ42. The reported lower affinities of shorter Aβ peptides towards γ-secretase may, at least partially, explain their decreased inhibitory potencies (*30*).

Our analysis on cultured neurons and neuron-like cells showed that extracellularly applied human Aβ42, but not p3, resulted in the accumulation of γ-secretase substrates. This phenotype is consistent with an Aβ42-induced inhibitory effect on γ-secretase activity, but could also be a consequence of alterations in cellular mechanisms that define protein (substrate) steady-state levels. However, Aβ42 treatment did not induce significant cellular toxicity, as determined by two independent readouts, nor did it promote APP-CTF generation, as indicated by the analysis of total, soluble APP ectodomain (precursor) levels. In addition, the assessment of the APP-CTF turnover by γ-secretase and/or general degradation mechanisms defined the inhibition of endogenous γ-secretases as the most important mechanism contributing to the accumulation of APP-CTF in living neurons treated with human Aβ42.

Intriguingly, despite their common inhibitory effects in cell-free conditions, only human Aβ42 and no other Aβ peptides induced the accumulation of unprocessed APP-CTFs in cells. Reports showing that Aβ conformation affects both cellular internalization and neurotoxicity (*74*) motivated us to test the possibility that Aβ42, unlike other Aβ peptides, acquires specific conformations that promote its endocytosis and/or inhibit its intracellular degradation. This would result in the intracellular concentration of this peptide at sites where γ-secretases process their substrates. Earlier reports of the selective uptake of Aβ42, relative to Aβ40 (*47, 75–77*), facilitating concentration of Aβ42 in spatially restricted endosomal compartments, would support this postulation. Our data showing marked increases in the intracellular levels of Aβ42, but neither Aβ40 nor Aβ43, add to the evidence for peptide-specific differences, and suggest that the observed increases mediate the inhibition of γ-secretases. Whether they are due to changes in Aβ42 cellular uptake, degradation rate or both requires further study.

Evidence that Aβ42 exerts a general inhibition on γ-secretase activity is provided by data showing that this peptide also reduced the processing of non-APP γ-secretase substrates, including NOTCH, ERBB4, neurexin, pan-cadherin and p75-based substrates. Considering the poor sequence homology between distinct γ-secretase substrates and the general Aβ-mediated inhibitory effect, we reason that the inhibitory action of Aβ42 involves its interaction with the protease, rather than with the substrate.

Our analysis also showed that Aβ42-induced inhibition of γ-secretases alters downstream signaling events. Treatment of BFCNs (or PC12 cells) with human Aβ42, but not with p3, compromised the processing of p75, increased p75-CTF levels and, in the absence of TrkA, induced apoptotic cell death. These observations recapitulated reported findings showing that GSI treatment of BFCNs triggers p75-driven cell death under similar conditions (*44*). It is noteworthy that these studies also showed that apoptosis in BFCNs subjected to GSI and TrkA signaling inhibitor treatments was rescued in p75 KO BFCNs, demonstrating that cell death is p75-dependent (*44*). The finding that Aβ42 treatment mimics GSI-driven p75-dependent apoptosis in BFCNs supports the inhibitory role of Aβ42 and links it to dysregulation of downstream signaling cascades. Our findings may also provide mechanistic bases for previous observations showing that injection of Aβ42 into the hippocampus of adult mice resulted in the degeneration of choline acetyltransferase (ChAT)-positive forebrain neurons in wildtype, but not p75-NTR-deficient mice, and revealing a correlation between Aβ42-driven neuronal death and the accumulation of p75-CTFs (*78, 79*).

Aβ endocytic uptake has been shown to concentrate Aβ42 to μM levels in the endolysosomal compartments (*47, 49*), where the high peptide concentration and low pH are proposed to facilitate Aβ aggregation and toxicity. We speculate that this mechanism could in addition promote the inhibition of γ-secretases. Given the reported role for APP_C99_ in inducing dysregulation of the endolysosomal network, the potential Aβ42-mediated inhibition of γ-secretase processing of APP-CTFs could drive dysregulation of endolysosomal function leading to changes in synaptic and axonal signaling (*41–43, 80–83*). Equally intriguing is the possibility that the general inhibition of γ-secretase substrates by Aβ42 could contribute to neuroinflammation by modifying microglia biology (*84*) and neurodegeneration, as reported previously for the genetic inactivation of these enzymes (*8–11*).

From a mechanistic standpoint, the competitive nature of the Aβ42-mediated inhibition implies that it is partial, reversible, and regulated by the relative concentrations of the Aβ42 peptide (inhibitor) and the endogenous substrates **(Figure 10C and 10D)**. The model that we put forward is that cellular uptake, as well as endosomal production of Aβ, result in increased intracellular concentration of Aβ42, facilitating γ-secretase inhibition and leading to the buildup of APP-CTFs (and γ-secretase substrates in general). As Aβ42 levels fall, the augmented concentration of substrates shifts the equilibrium towards their processing and subsequent Aβ production. As Aβ42 levels rise again, the equilibrium is shifted back towards inhibition. This cyclic inhibitory mechanism will translate into pulses of (partial) γ-secretase inhibition, which will alter γ-secretase mediated-signaling (arising from increased CTF levels at the membrane or decreased release of soluble intracellular domains from substrates). These alterations may affect the dynamics of systems oscillating in the brain, such as NOTCH signaling, implicated in memory formation, and potentially others (related to e.g. cadherins, p75 or neuregulins). It is worth noting that oscillations in γ-secretase activity induced by treatment with a γ-secretase inhibitor semagacestat have been proposed to have contributed to the cognitive alterations observed in semagacestat treated patients in the failed Phase-3 IDENTITY clinical trial (*7*) and that semagacestat, like Aβ42, acts as a high affinity competitor of substrates (*85*).

The convergence of Aβ42 and tau at the synapse has been proposed to underlie synaptic dysfunction in AD (*86–89*), and recent assessment of APP-CTF levels in synaptosome-enriched fractions from healthy control, SAD and FAD brains (temporal cortices) has shown that APP fragments concentrate at higher levels in the synapse in AD-affected than in control individuals (*90*). Our analysis adds that endogenous Aβ42 concentrates in synaptosomes derived from end-stage AD brains to reach ∼10 nM, a concentration that in CM from human neurons inhibits γ-secretase in PC12 cells (**Figure 7**). Furthermore, the restricted localization of Aβ in endolysosomal vesicles, within synaptosomes, likely increases the local peptide concentration to levels that inhibit γ-secretase-mediated processing of substrates in this compartment. In addition, we argue that the deposition of Aβ42 in plaques may be preceded by a critical increase in the levels of Aβ present in endosomes and the cyclical inhibition of γ-secretase activity that we propose. Under this view, reductions in γ-secretase activity may be a (transient) downstream consequence of increases in Aβ due to failed clearance, as represented by plaque deposition, contributing to AD pathogenesis.

The Aβ-mediated inhibition of γ-secretase may also help to explain the intriguing accumulation of APP-CTFs in the heterozygous FAD brain (*91*). In this regard, the direct quantification of γ-secretase activity in detergent resistant fractions prepared from post-mortem brain samples of healthy controls and FAD-linked mutation carriers revealed similar overall γ-secretase activity levels, indicating that the wildtype (PSEN1 and PSEN2) γ-secretase complexes rescue any potential mutation-driven deficits in the processing of APP (*92*). Yet APP-CTFs have been reported to accumulate in the FAD brain (*90, 91*) and the accumulation of APP-CTFs appear to correlate with Aβ levels at the synapse. The inhibition of γ-secretase by Aβ42 could resolve the apparent conflict. Indeed, our data could reconcile these two seemingly exclusive hypotheses on the effects of FAD mutations in *PSEN1* on the development of AD by noting that: 1) there is mutation-driven enhanced generation of Aβ42 within the endolysosomal network; 2) that through both endosomal production and endocytosis Aβ42 increases to a level within the endolysosomal network sufficient to inhibit the γ-secretase complex; and 3) that in the case of FAD mutations the isolation of the γ-secretase releases Aβ42, thus restoring wildtype enzyme activity (*28, 93*). Thus, increased levels of endolysosomal Aβ42 with concurrent inhibition of γ-secretase may be responsible, at least in part, for the apparent γ-secretase loss-of-function phenotypes.

Collectively, our data raise the intriguing possibility that increases in Aβ42 in the AD brain, and in particular in the endolysosomal compartment, facilitate the establishment of an Aβ-driven inhibitory mechanism that contributes to neurotoxicity by impairing critical γ-secretase signaling functions. By mechanistically connecting elevated Aβ42 levels with the accumulation of multiple γ-secretase substrates, our observations integrate disparate views as to which pathways lead to neurodegeneration and offer a novel conceptual framework for investigating the molecular and cellular bases of AD pathogenesis.

## MATERIALS AND METHODS

### Chemicals, peptides and antibodies

Aβ peptides were purchased from rPeptide, resuspended in DMSO at 500 μM, aliquoted into single use 10 μl aliquots and stored at -80°C. For Aβ42 the following lots were used: 4261242T, 06021342T and 02092242T. Γ-secretase inhibitors (Inhibitor X (InhX, L-685,458), DAPT and compound E (CE)) were purchased from Bioconnect, Sigma Aldrich and Millipore, respectively. TrkA inhibitor K252α, cycloheximide and Bafilomycin A1 were purchased from Sigma Aldrich. The following antibodies were used: mouse anti-FLAG M2 (Sigma Aldrich, F3165), rabbit anti-ADAM10 antibody (EPR5622, Abcam, ab124695), rabbit anti-APP (gift from Prof. Wim Annaert (B63)), rabbit anti-APP (Y188, Abcam, ab32136), mouse anti-APP (22C11, Thermo Fisher Scientific, 14-9749-82), rabbit anti-BACE1 (EPR19523, Abcam, ab183612), rabbit anti-pan-cadherin (Thermo Fisher Scientific, 71-7100), anti-p75 NTR (Millipore, 07-476), anti-Aβ (clone 4G8, Biolegend), anti-choline acetyltransferase (Millipore, AB144P), anti-cleaved caspase-3 (Cell Signaling, 9661S), HRP-conjugated goat anti-rabbit (BioRad), Alexa Fluor 790-conjugated goat anti-mouse (Thermo Fisher Scientific), Cy3-conjugated donkey anti-rabbit (Jackson ImmunoResearch Laboratories), Alexa Fluor Plus 488-conjugated donkey anti-mouse (Thermo Fisher Scientific), biotinylated rabbit anti-goat (Jackson ImmunoResearch Laboratories) and Cy2-conjugated streptavidin (Jackson Immunoresearch Laboratories).

### AAV production

Preparation of the AAV-hSyn1-C99 Y-T biosensor was performed as described previously (*53*). Briefly, the cDNA of C99 Y-T probe was subcloned into a pAAV2/8 vector containing human synapsin 1 promoter and WPRE sequences (*94*). The packaging into viruses was performed at University of Pennsylvania Gene Therapy Program vector core (Philadelphia, PA). The virus titer was 4.95E+13 GC/ml.

### Expression and purification of γ secretase and γ-secretase substrates

High Five insect cells were transduced with baculoviruses carrying cDNA encoding all γ-secretase subunits (wildtype PSEN1 or PSEN2, NCSTN, APH1A or APH1B, PEN2) and 72 hours later the cells were collected for protein purification, as described before (*30*).

Recombinant γ-secretase substrates were expressed and purified using mammalian cell expression system or baculovirus-mediated expression system in High Five insect cells as before (*30, 44, 95*). COS cells were transfected with pSG5 plasmid encoding NOTCH1- or P75-based γ-secretase substrate, tagged at the C-terminus with 3xFLAG. High Five insect cells were transduced with baculoviruses carrying cDNA encoding APP_C99_, ERBB4ΔECT and neurexinΔECT-tagged with 3xFLAG-prescission protease (PPS) cleavage site-GFP tandem at the C-terminus. The purity of the protein was analyzed by SDS-PAGE and Coomassie staining (InstantBlue Protein Stain, Expedeon).

### Preparation of detergent resistant membranes

CHAPSO detergent resistant membranes (DRMs) were prepared from High Five insect cells overexpressing wildtype γ-secretase complexes containing NCSTN, APH1B, PSEN1 and PEN2 subunits, as reported before (*30*).

### Cell-free γ-secretase activity assays

Purified γ-secretases (∼25 nM) were mixed with respective, purified 3xFLAG-tagged γ-secretase substrates (0.4 or 1.2 μM) in 150 mM NaCl, 25 mM PIPES, 0.25% CHAPSO, 0.03% DDM, 0.1% phosphatidylcholine buffer and the reactions were incubated for 40 minutes at 37°C. Non-incubated reactions or reactions supplemented with 10 μM InhX served as negative controls. In the experiments testing for the reversibility of the inhibition, γ-secretases were first immobilized on sepharose beads covalently coupled to anti-γ-secretase nanobody. Following the first reaction containing Aβ or GSI, the beads were washed 3x for a total duration of 30 minutes with 150 mM NaCl, 25 mM PIPES, 0.25% CHAPSO buffer and a fresh substrate was added. For γ-secretase activity assays in membrane-like conditions (25 μl total reaction volume), 6.25 μl DRMs, re-suspended in 20 mM PIPES, 250 mM sucrose, 1 M EGTA, pH7 at the concentration of 1 μg/μl, were mixed with 6.25 μl APP_C99_-3xFLAG or Aβ1-42 substrate in 150 mM NaCl, 25 mM PIPES, 0.03% DDM, 2.5 μl 10xMBS and H_2_O supplemented with DDM and CHAPSO, to achieve final concentration of the detergents 0.03% and 0.1%, respectively, and final concentration of the substrates 1.5 μM and 10 μM, respectively. The reaction mixes were incubated for 40 minutes at 37°C. Reactions supplemented with 10 μM InhX served as negative controls.

### Analysis of ICD generation in cell-free γ-secretase activity assays

*De novo* ICD generation was determined by western blotting. Briefly, the reaction mixtures were subjected to methanol:chloroform (1:2 v/v) extraction to remove the excess of unprocessed substrates, and the upper fractions (containing mainly the generated ICDs) subjected to SDS-PAGE. Otherwise, the high levels of substrate could preclude quantitative analysis of ICDs **(Supplementary figure 1).**

ICD-3xFLAG were detected by western blotting with anti-FLAG, followed by anti-mouse Alexa Fluor 790 conjugated antibodies. Fluorescent signals were developed using Typhoon (GE Healthcare). The intensity of the bands was quantified by densitometry using ImageQuant software.

Alternatively, MALDI-TOF MS was applied to determine the relative *de novo* ICD-3xFLAG levels generated in cell-free conditions, as reported before (*30*). The samples were spiked with Aβ1-28 as an internal standard. The mass spectra were acquired on a RapiFleX TOF mass spectrometer (Bruker Daltonics) equipped with a 10 kHz Smartbeam™ laser using the AutoXecute function of the FlexControl 4.2 acquisition software.

### Analysis of γ-secretase substrate proteolysis in cultured cells using secreted HiBiT-Aβ or -Aβ-like peptide levels as a proxy for the global γ-secretase endopeptidase activity

HEK293 stably expressing APP-CTF (C99) or a NOTCH1-based substrate (similar in size as APP-C99) both N-terminally tagged with the HiBiT tag were plated at the density of 10000 cells per 96-well, and 24h after plating treated with Aβ or p3 peptides diluted in OPTIMEM (Thermo Fisher Scientific) supplemented with 5% FBS (Gibco). Conditioned media was collected and subjected to analysis using Nano-Glo® HiBiT Extracellular Detection System (Promega). Briefly, 50 µl of the medium was mixed with 50 µl of the reaction mixture containing LgBiT Protein (1:100) and Nano-Glo HiBiT Extracellular Substrate (1:50) in Nano-Glo HiBiT Extracellular Buffer, and the reaction was incubated for 10 minutes at room temperature. Luminescence signal corresponding to the amount of the extracellular HiBiT-Aβ or -Aβ-like peptides was measured using victor plate reader with default luminescence measurement settings.

### Analysis of γ-secretase substrate levels in cultured cells

SH-SY5Y, PC12 and PC12nnr5 (*59*) cell lines were cultured in Dulbecco’s Modified Eagle Medium (DMEM)/F-12 (Thermo Fisher Scientific) supplemented with 10% fetal bovine serum (FBS) (Gibco) at 37°C, 5% CO_2_. ReNCell VM cells were cultured in Corning® Matrigel® hESC-Qualified Matrix, LDEV-free-coated flasks in DMEM/F-12 medium supplemented with B27 (Thermo Fisher Scientific), 2 μg/ml heparin (Stem Cell Technologies), 20 ng/ml EGF-1 (Cell Guidance) and 25 ng/ml FGF (Cell Guidance). Human iPSC line was derived from a cognitively unaffected male individual, previously established and characterized (CVB) (RRID: CVCL_1N86, GM25430) (*96*). The two neuronal progenitor cell (NPC) lines – CV4a (derived from CVB iPSC line that carries a wild type APP, herein termed as WT) and APPSwe (homozygous for APP Swedish mutations that were genome-edited from CVB parental iPSC line) were characterized and reported (*97, 98*). NPCs were plated on 20 μg/ml poly-L-ornithine (Sigma Aldrich) and 5 μg/ml laminin (Thermo Fisher Scientific)-coated plates in NPC media: DMEM/F12/Glutamax supplemented with N2, B27, penicillin and streptomycin (Thermo Fisher Scientific) and 20 ng/ml FGF-2 (Millipore). For neuronal differentiation, confluent NPCs were cultured in NPC media without FGF-2 for three weeks. All cells were maintained in a humidified incubator at 37°C, 5% CO_2_, and regularly tested for mycoplasma infection.

For the cellular assays, cells were plated into 6-well plates at the density of 300000 cells per well and 24h after plating treated with Aβ or p3 peptides diluted in OPTIMEM (Thermo Fisher Scientific) supplemented with 5% FBS (Gibco). In the experiments set to analyze sAPP in the conditioned medium, FBS was replaced with 1% knock-out serum replacement (KOSR) (Thermo Fisher Scientific) due to the interference of the antibodies present in the serum with western blotting. After 8h, the medium was refreshed, using peptide supplemented media, and the cultures were incubated at 37°C for 16h. Cell lysates were prepared in radioimmunoprecipitation (RIPA) buffer (25 mM Tris, 150 mM NaCl, 1% Nonidet P-40, 0.5% sodium deoxycholate, 0.1% SDS, pH 7.4) supplemented with protease inhibitors. Equal amounts of protein or conditioned medium were subjected to SDS-PAGE on 4-12% Bis-Tris gels in MES or MOPS buffer or 4-16 % Tris-Tricine gels (*99*) and western blot analysis. Chemiluminescent signals were developed using Fujifilm LAS-3000 Imager or ChemiDoc XRS+ imaging apparatus (BioRad). The optical density of the bands was quantified using ImageQuant or Image J software.

### Cytotoxicity assays

Cytotoxicity was assessed using LDH-Glo Cytotoxicity Assay (Promega), which is a bioluminescent plate-based assay for quantifying LDH release into the culture medium upon plasma membrane damage. The assay was performed following the manufacturer recommendation. Briefly, culture medium was collected upon respective treatment and diluted 1 in 100 in LDH storage buffer (200mM Tris-HCl (pH 7.3), 10% glycerol, 1% BSA). Medium from cells treated for 15 min with 2% Triton X-100 was used as a maximum LDH release control. 12.5 µl of diluted medium was mixed with 12.5 µl LDH detection enzyme mix supplemented with reductase substrate in 384-well plate. The reactions were incubated for 1h and luminescence read using Promega GloMax Discoverer.

Cytotoxicity was also assessed using CellTiter-Glo® 2.0 Assay (Promega), which is a homogeneous method used to determine the number of viable cells based on quantitation of the ATP, an indicator of metabolically active cells. Briefly, cells were plate in 96-well plate in 100 µl medium, treated following the treatment regime, mixed with CellTiter-Glo reagent and luminescence was read using Promega GloMax Discoverer.

### Isolation of biologically derived human Aβ and cellular assay for γ-secretase-mediated proteolysis

The conditioned media from the WT (WT-CM) and APPSwe (SWE-CM) human neurons were used as a biological source of Aβ. Following three weeks of differentiation, neurons cultured on 10 cm diameter dishes were incubated in NPC media without FGF-2 for five days. After five days, conditioned media were collected in a sterile atmosphere. Immunodepletion of Aβ from APPSwe medium was done following a modified protocol (*100*). Briefly, 5 ml of SWE-CM were incubated with 3D6 antibody (1 µg/ml), 15 µl protein A and 15 µl protein G resin for 3h at 4°C with gentle shaking on a nutator (*101*). The supernatant was collected after centrifuging at 3500×g and used as immunodepleted media (delSWE-CM) for cell culture experiments. To test for the APP-CTF accumulation in the presence of different conditioned media, PC12 cells, cultured on 6-well plates, were treated with WT-CM, SWE-CM or delSWE-CM for 24h along with regular feeding media (control). Following incubation, cells were harvested and processed to detect APP-FL and APP-CTF.

### Spectral FRET analysis of γ-secretase activity in living neurons

Primary neuronal cultures were obtained from cerebral cortex of mouse embryos at gestation day 14–16 (Charles River Laboratories). The neurons were dissociated using Papain Dissociation System (Worthington Biochemical Corporation, Lakewood, NJ) and maintained for 13-15 days *in vitro* (DIV) in neurobasal medium containing 2% B27, 1% GlutaMAX Supplement and 1% penicillin/streptomycin (Thermo Fisher Scientific). A laser at 405 nm wavelength was used to excite TurquoiseGL in the C99 Y-T biosensor (*53*). The emitted fluorescence from the donors (TurquoiseGL) and the acceptors (YPet) was detected at 470±10nm and 530±10nm, respectively, by the TruSpectral detectors on an Olympus FV3000RS confocal microscope equipped with CO_2_/heating units. x10/0.25 objective was used for the imaging. Average pixel fluorescence intensity for the cell body after subtraction of the background fluorescence was measured using Image J. The emission intensity of YPet over that of TurquoiseGL (Y/T) ratio was used as a readout of the FRET efficiency, which reflects the relative proximity between the donor and the acceptor. Pseudo-colored images were generated in MATLAB (MathWorks, Natick, MA). The neuronal preparation procedure followed the NIH guidelines for the use of animals in experiments and was approved by the Massachusetts General Hospital Animal Care and Use Committee (2003N000243).

### Primary cultures of basal forebrain cholinergic neurons

Animals of CD1 genetic background were housed in an animal care facility with a 12-hour dark/light cycle and had free access to food and water. All experiments were performed according to the animal care guidelines of the European Community Council (86 ⁄ 609 ⁄ EEC) and to Spanish regulations (RD1201 ⁄ 2005), following protocols approved by the ethics committees of the Consejo Superior Investigaciones Científicas (CSIC). Septal areas were dissected from CD1 E17-18 embryos in chilled Hanks’ balanced salt solution (HBSS, 137 mM NaCl, 5.4 mM KCl, 0.17 mM Na_2_HPO_4_, 0.22 mM KH_2_PO_4_, 9.9 mM HEPES, 8.3 mM glucose and 11 mM sucrose). The septa were pooled and digested with 1 ml of 0.25% trypsin (GE Healthcare) and 0.5 ml of 100 kU DNase I (GE healthcare) for 10 minutes at 37°C. The tissue was further dissociated by aspiration with progressively narrower tips in neurobasal medium (Thermo Fisher Scientific) supplemented with 4% bovine serum albumin (BSA), 2% B27 (Thermo Fisher Scientific), 1% L-glutamine (Thermo Fisher Scientific) and 0.5% penicillin/streptomycin (GE Healthcare) (NB/B27). After tissue dissociation, equal volume of 4% BSA was added and samples centrifuged at 300xg for 5 minutes at 4°C. The supernatant was discarded and the cell pellet resuspended in 5 ml of 0.2% BSA. The suspension was filtered through 40 µm nylon filter (Sysmex), cells counted in Neubauer chamber, centrifuged again and resuspended in NB/B27. Cells were seeded in 24-well plates (2×10^5^ cells/well) containing 12 mm diameter pre-coated coverslips (VWR) for immunocytochemical analysis or in pre-coated p-100 plates for western blotting. The surfaces were coated with 50 µg/ml poly-D-lysine (Sigma Aldrich) overnight at 4°C and 5 µg/ml laminin (Sigma Aldrich) for 2h at 37°C. The next day, half of the medium was exchanged and the concentration of B27 reduced to 0.2%. The medium was supplemented with 2 µM of the antimitotic agent (1-β-D-Arabinofuranosylcytosine (Sigma Aldrich)) and 100 ng/ml NGF (Alomone labs). Neurons were cultured until DIV11. The respective treatments were applied at DIV8 and the culture continued for 72h until DIV11.

### Immunocytochemical analysis of the cell death rate

For immunocytochemistry (ICC) cells were fixed in 2% paraformaldehyde and permeabilized with 0.1% Triton X-100. After that they were incubated with 0.5% SDS for 5 minutes for antigen retrieval. Non-specific binding of antibodies was blocked using 2% BSA, 0.1% Triton X-100 in 0.1 M phosphate buffer (PB). The cells were incubated overnight with anti-choline acetyltransferase (Millipore) (only for BFCN) and anti-cleaved caspase-3 (Cell Signaling) antibodies at 4°C. The next day the samples were incubated with corresponding secondary antibodies: Cy3-conjugated donkey anti-rabbit, biotinylated rabbit anti-goat, and Cy2-conjugated streptavidin (all purchased from Jackson Immunoresearch Laboratories). Nuclei were stained with DAPI and samples mounted on glass slides. For quantitative analysis, for BFCNs total number of choline acetyltransferase positive neurons and the number of double positive neurons for choline acetyltransferase and cleaved caspase-3 was determined, while for PC12 a fraction of cleaved caspase 3 positive cells among all cells (determined by DAPI staining) was determined.

### Immunocytochemical analysis of Aβ internalization

PC12 cells were plated on glass coverslips and treated for 24h with respective Aβ peptides at 1 μM. The cells were fixed with 4% paraformaldehyde, and permeabilized with 0.5% Triton X-100 in TBST (0.2% Tween 20 in TBS). Blocking was performed by 1h incubation in solution containing 1% bovine serum albumin, 0.1% gelatin, 300 mM glycine and 4% normal donkey serum in TBST. Primary antibodies were diluted in the blocking solution and incubated overnight. This was followed by incubation with respective Alexa Fluor Plus 488 conjugated secondary antibody and Alexa Fluor Plus 555 conjugated phalloidin. The samples were then incubated with nuclear stain DAPI for 5 min and mounted using Prolong glass. Images were acquired at 63x magnification using Zeiss LSM 900 confocal microscope at VIB Bioimaging Core.

### Measurement of Aβ42 levels in synaptosomes from post-mortem human cortical tissue

The *post mortem* human brain tissues from the frontal cortex were obtained from the University of California San Diego–Alzheimer’s Disease Research Center (UCSD-ADRC; La Jolla, CA, USA), and University of California Irvine–Alzheimer’s Disease Research Center (UCI-ADRC) brain tissue repository (Irvine, CA, USA; **Supplementary table 2**), and stored at -80°C prior to use. Synaptosomes from the cortices were isolated following published protocol using a discontinuous Percoll gradient (Sigma Aldrich, USA) (*62, 63*) with some modifications. All the steps were performed on ice or at 4°C unless mentioned otherwise. Briefly, 1200 mg of tissue was homogenized using a handhold Dounce homogenizer in 5 volumes of homogenization buffer (HB: 0.32 M sucrose, 5 mM magnesium chloride, 5 mM Tris-Cl, pH 7.4 and EDTA-free protease inhibitor cocktail). After centrifuging at 1000xg for 10 minutes, post-nuclear supernatant (PNS) was collected. The PNS was further centrifuged at 10800xg for 10 minutes to isolate the pellet fractions (P2, crude synaptosome). The P2 fraction was resuspended in HB and loaded on top of a three-step (3%, 10% and 23%) Percoll gradient, and centrifuged at 14500xg for 12 minutes at 4°C in a Fiberlite F21-8 x 50y Fixed-Angle Rotor (Thermo Fisher, USA). The synaptosome-rich interface between 10% and 23% Percoll layers was collected and resuspended in 30 volumes of HB. The diluted material was centrifuged at 18500xg for 30 minutes at 4°C, and the synaptosome-enriched pellet was resuspended in 500 μl of HB. To measure the concentration of Aβ42, 25 μl of synaptosome suspension was centrifuged at 18500xg for 10 minutes at 4°C. Supernatant was discarded and the volume of synaptosome pellet was estimated by adding minimum volume of liquid (2 μl of PBS) to the synaptosome pellet. Synaptosomes were dissolved in 23 μl RIPA buffer (25 mM Tris-HCl pH 7.5, 150mM NaCl, 1% NP-40, 1% sodium deoxycholate, 0.1% SDS) containing Halt Protease and Phosphatase Inhibitor Cocktail (Thermo Fisher Scientific). After centrifugation for 15 minutes at 13,800xg at 4°C, the supernatant was collected for MSD-ELISA (Meso Scale Diagnostics, USA). Concentrations of Aβ were measured using the VPLEXAβ Peptide Panel (4G8: MesoScaleDiscovery) following the manufacturer’s Instructions. Data were obtained using the MESO QuickPlex SQ 120 and analyzed using DISCOVERY WORKBENCH 3.0 (Meso Scale Diagnostics). The pg concentration of Aβ in 25 μL was converted to pg/mL and eventually to nM values using the volume of synaptosomes.

### Measurement of APP-CTF levels in peptide-treated synaptosomes from mouse cortical tissue

The cortical tissues from two-months old wild-type mice (N=3) were collected and stored at -80°C. Synaptosomes from the 60-70 mg of cortices were isolated following the protocol described above. The synaptosome-rich interface between 10% and 23% Percoll layers was collected and resuspended in 30 volumes of HB. The diluted material was centrifuged at 18500xg for 30 minutes at 4°C, and the synaptosome-enriched pellet was resuspended in HB supplemented with 10 mM glucose. 10-15 μg of synaptosome was incubated with Aβ42 peptide at 2.5 μM final concentration at 37°C for 18h. DMSO was used as a vehicle control. One synaptosomal sample was treated with 200 nM of Compound E. Following incubation, samples were resolved on SDS-PAGE and western blotting was performed using anti-APP Y188 and anti-GAPDH antibodies. All densitometric analyses were performed using NIH ImageJ software. The animal experiments were approved by the Institutional Animal Care and Use Committee of the University of California San Diego.

### Statistics

Statistical analysis was performed using Excel, GraphPad Prism, R 4.2.2. and R Studio software. The following R packages were used for the analysis: readxl, ggplot2, plyr, dplyr, DescTools, gridExtra and reshape2 (*102–104*). P<0.05 was considered as a predetermined threshold for statistical significance. One-way or two-way ANOVA, or Kruskal-Wallis test followed by Dunnett’s, Tukey’s or Dunn multiple comparison test or unpaired Student’s t-test were used, as described in the legends.

## ACKNOWLEDGEMENTS

This work was supported by the Cure Alzheimer’s Fund (WCM-LCG), the Research Foundation Flanders (FWO, Research project G0B2519N) (LCG), the DH Chen Foundation (WCM), NIH AG015379 (OB), NIH AG044486 (OB), NIH R01AG055523 (WCM) and Bundesministerium für Bildung und Forschung (BMBF) grant M^2^OGA within the Partnership for Innovation in Health Industry, M^2^Aind (CH). MV acknowledges the Spanish Ministry of Science and Innovation (grant PID2021-127600NB-I00). We would like to thank VIB Bioimaging Core for support with confocal imaging, VIB Flow Core (Dr Jochen Lamote and Gonzalo Delgado Martinez) for support with FACS and Dr Laetitia Miguel for support in the culture of ReNcell VM cells. We thank all the Mobley and Chávez-Gutiérrez lab members for fruitful discussions.

## Supplementary figures

**Supplementary figure 1.**
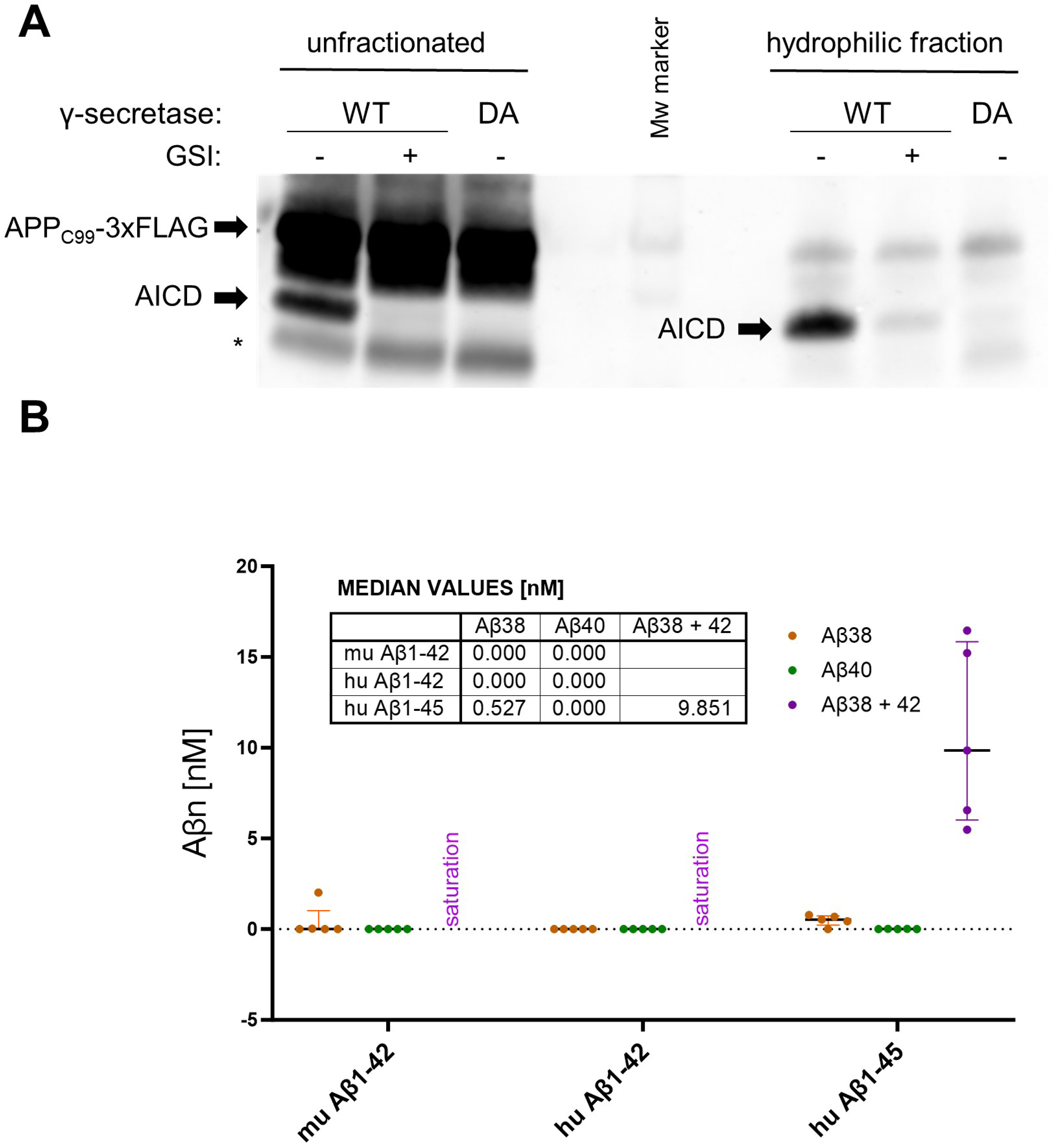
Cell-free γ-secretase activity assay. (A) Analysis of AICD-3xFLAG levels that are *de novo* generated by purified γ-secretase from the APP_C99_-3xFLAG substrate is shown. Γ-secretase inhibitor (GSI) and catalytically inactive γ-secretase mutant (DA) were used as negative controls. The left side of the blot presents analysis of unfractionated γ-secretase activity assay samples, while the hydrophilic fraction obtained through methanol:chloroform extraction in presented on the right side. This protein extraction method allows quantification of AICDs without interference of the signal coming from the excess of APP_C99_-3xFLAG substrate present in the assays. * marks a hydrophobic contaminant. (B) The graph presents ELISA quantification of Aβ1-38, 1-40 and 1-42 peptides generated in detergent-based γ-secretase activity assays using either human Aβ1-42, murine Aβ1-42 or human Aβ1-45 at 10 μM as substrates. The data are presented as median ± IQR, N=5.

**Supplementary figure 2.**
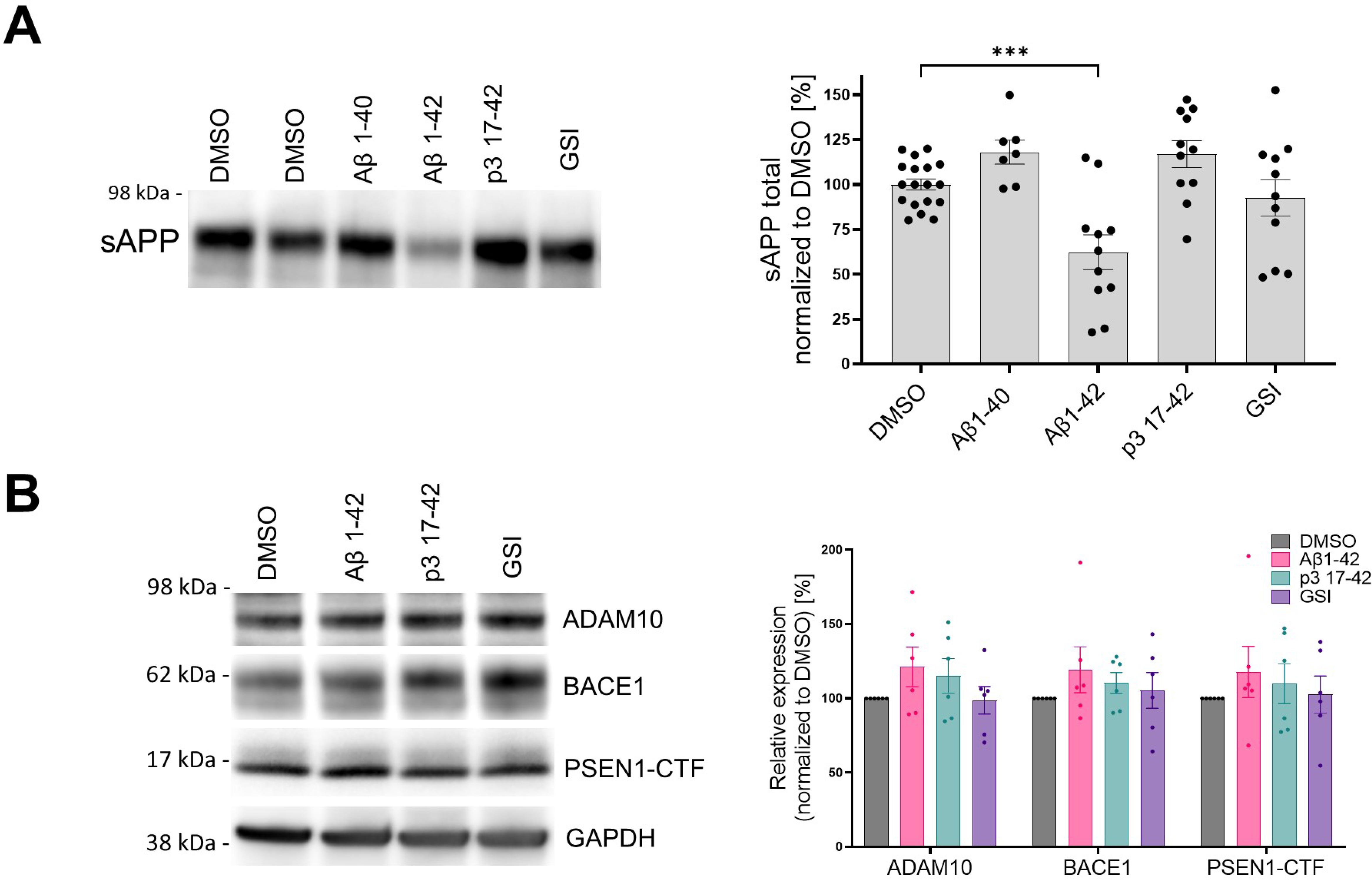
Sheddase activation is not behind the APP-CTF accumulation in cells. (A) The western blots present soluble APP (sAPP) detected in conditioned medium collected from SH-SY5Y cells treated for 24h with respective peptides at 1 μM concentration, GSI (Inh X, 2 μM) or vehicle (DMSO). The sAPP levels were calculated from the integrated density of the corresponding western blot bands. The data are shown as mean ± SEM, N=7-18. The statistics were calculated using one-way ANOVA and multiple comparison Dunnett’s test, with DMSO set as a reference. ***p<0.001. (B) The western blots present ADAM10, BACE1, PSEN1-CTF and GAPDH detected in total lysates prepared from SH-SY5Y cells treated for 24h with respective peptides at 1 μM concentration, GSI (InhX, 2 μM) or vehicle (DMSO). The relative expression levels of the proteases were calculated from the integrated density of the corresponding western blot bands and normalized to housekeeping protein levels (GAPDH). The data are shown as mean ± SEM, N=6, with DMSO considered as 100%. The statistics were calculated using two-way ANOVA and multiple comparison Dunnett’s test, with DMSO set as a reference. No statistically significant differences were detected between the tested conditions.

**Supplementary figure 3.**
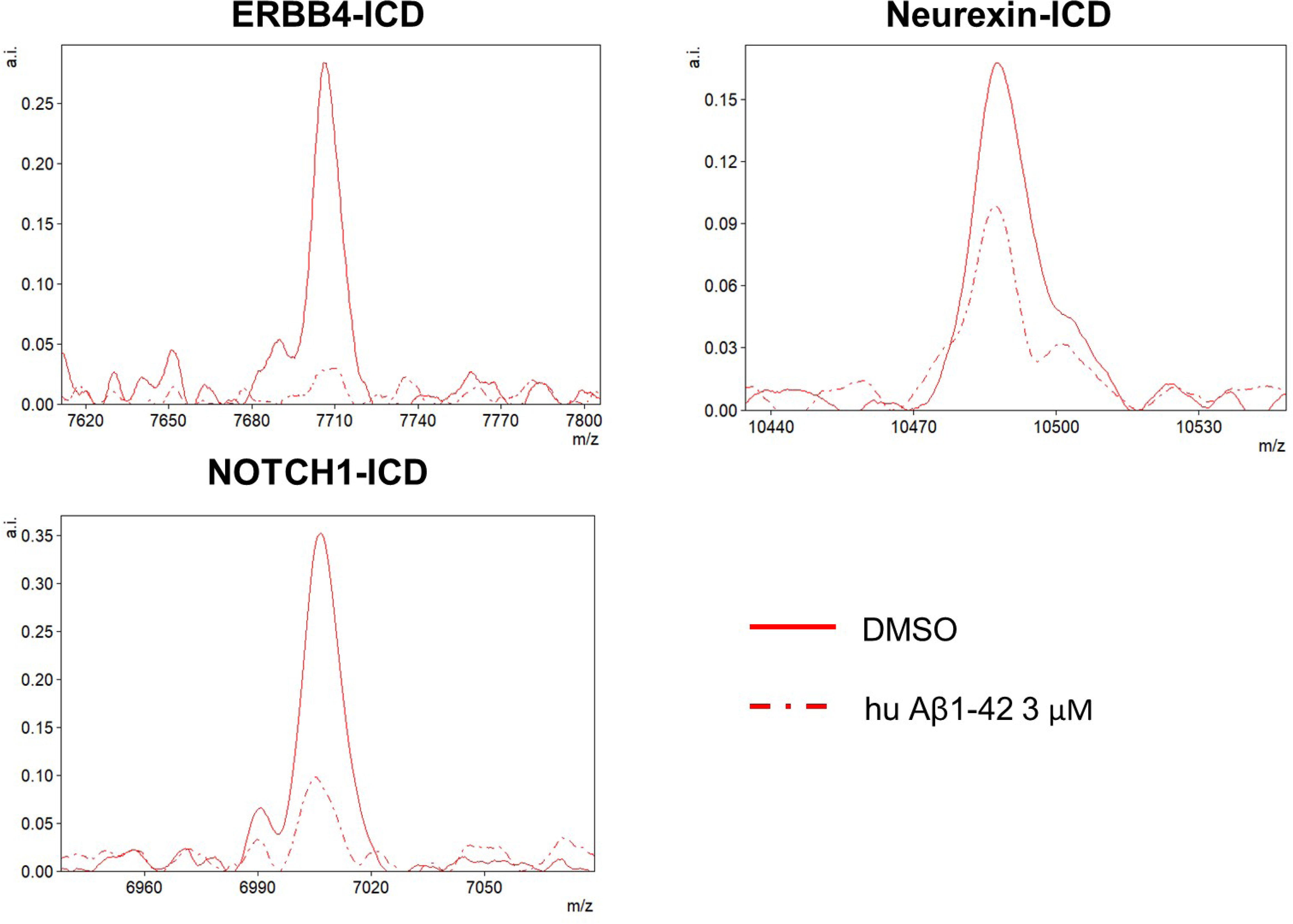
Mass spectrometry analysis confirms the inhibitory action of human Aβ1-42 peptides. The chromatograms present the generation of ICDs from three different substrates in *in vitro* cell-free γ-secretase activity assays, analyzed by mass spectrometry. Solid red – vehicle; dashed red – 3 μM human Aβ1-42.

## Supplementary Tables

**Supplementary Table 1.** Summary of IC50s in cell-free detergent-based γ-secretase activity assays for selected Aβ peptides.

|  |  | IC<sub>50</sub> [nM] |  |
| --- | --- | --- | --- |
| | | Substrate: 0.4 μM | Substrate: 1.2 μM |
| AICD | Aβ1-42 | 1259 (N=9) | 3565 (N=5) |
| | Aβ11-42 | 2218 (N=3) | 4685 (N=4) |
|  | p3 17-42 | N/A | N/A |
| | murine Aβ1-42 | N/A | N/A |
| NICD | Aβ1-42 | 609 (N=3) | 1564 (N=3) |

**Supplementary table 2.** Demographics of the human post-mortem brain tissue.

| Origin | Category | Case # | Sex | Age | PMI (h) | Clinical diagnosis | Additional comments | Braak stage | APOEεX/<br>εX status<br>(X = 2, 3,<br>and/or<br>4) |
| --- | --- | --- | --- | --- | --- | --- | --- | --- | --- |
| ADRC-UCI | ND | 19-13 | F | 87 | 3.83 | Normal |  | III | 2/3 |
| ADRC-UCI | ND | 25-01 | M | 83 | 1.8 | Normal |  | II | 3/3 |
| ADRC-UCSD | ND | 5687 | M | 84 | 8 | Normal |  | I | 3/3 |
| ADRC-UCSD | ND | 5783 | M | 84 | 36 | Normal |  | II | 2/3 |
| ADRC-UCSD | ND | 5844 | F | 96 | 6 | Normal |  | II | 3/3 |
| ADRC-UCSD | ND | 5919 | F | 75 |  | Normal | Arteriosclerosis | I | 2/3 |
| ADRC-UCSD | AD | 5401 | M | 81 | 8 | Alzheimer's disease | Amyloid angiopathy | IV | 4/4 |
| ADRC-UCSD | AD | 5620 | M | 86 | 8 | Alzheimer's disease |  | IV | 3/3 |
| ADRC-UCSD | AD | 5835 | F | 85 | 30 | Alzheimer's disease | Lewy body disease-neocortical (diffuse) type | III | 3/3 |
| ADRC-UCSD | AD | 5851 | M | 71 | -4 | Alzheimer's disease | Lewy body disease-neocortical (diffuse) type | IV | 3/4 |
| ADRC-UCSD | AD | 5881 | F | 88 | -4 | Alzheimer's disease | Progressive supranuclear palsy, TDP-43 proteinopathy | III | 3/3 |
| ADRC-UCSD | AD | 5901 | F | 87 | -4 | Alzheimer's disease | Amyloid angiopathy, Lewy body pathology, amygdala predominant type | IV | 3/3 |

**Abbreviations:** ADRC, Alzheimer's Disease Research Center; UCI, University of California Irvine; UCSD, University of California San Diego; ND, Non-demented control; AD, Alzheimer's disease; PMI, post-mortem interval. Specimens and their diagnoses were provided by the brain banks at respective ADRCs.

## Notes

### Competing Interest Statement

KMZ, UD, TE, SL, MM, MCH, MLF, BO, CH, AB, DGM, DK, MV, OB, LCG declare no conflict of interests.
WM received grant or contract funding from the NIH, Ono Pharma Foundation, Cure Alzheimer Fund, DH Chen Foundation, AC Immune, Larry L Hillblom Foundation, Alzheimer Association, Annovis-Bio and BioSplice and the Michael J Fox Foundation. He consulted for Samumed and AC immune. He served on advisory boards for the Bluefield Project to Cure Frontotemporal Dementia, the Blythedale Burke Pediatric Neuroscience Research Collaboration, The Key, the National Down Syndrome Society, the American Neurological Association, the Sanford Health Lorraine Cross Award Committee, the NIH COBRE program at the University of Nebraska, the Dementia Aware Committee and the Dementia Committee for the State of California Health Services, and the San Diego Alzheimer Project. He was a member of a Pfizer Data Safety Monitoring Board. He serves as President of the T21 Research Society. Annovis Bio provided a gift to the WM lab and a test compound. WM received stock or stock options from Annovis-Bio, Alzheon, Curasen, Cortexyme, and Promis. He is co-inventor on UCSD Patents for Gamma-secretase Modulators. He received a travel reimbursement from AC Immune. He served as expert witness for Korein-Tillery, LLC.

### Summary of Updates

This is a revised version that presents new data.

## REFERENCES

1. L. Chávez-Gutiérrez, M. Szaruga, Mechanisms of neurodegeneration - Insights from familial Alzheimer’s disease. Semin Cell Dev Biol 105, 75–85 (2020).

2. D. J. Selkoe, J. Hardy, The amyloid hypothesis of Alzheimer’s disease at 25 years. EMBO Mol Med 8, 595–608 (2016).

3. A. Haapasalo, D. M. Kovacs, The many substrates of presenilin/γ-secretase. J Alzheimers Dis 25, 3–28 (2011).

4. G. Güner, S. F. Lichtenthaler, The substrate repertoire of γ-secretase/presenilin. Semin Cell Dev Biol 105, 27–42 (2020).

5. N. Jurisch-Yaksi, R. Sannerud, W. Annaert, A fast growing spectrum of biological functions of γ-secretase in development and disease. Biochim Biophys Acta 1828, 2815–2827 (2013).

6. C. M. Carroll, Y.-M. Li, Physiological and pathological roles of the γ-secretase complex. Brain Research Bulletin 126, 199–206 (2016).

7. R. S. Doody et al., A phase 3 trial of semagacestat for treatment of Alzheimer’s disease. N Engl J Med 369, 341–350 (2013).

8. H. Acx et al., Inactivation of γ-secretases leads to accumulation of substrates and non-Alzheimer neurodegeneration. EMBO Mol Med 9, 1088–1099 (2017).

9. M. Wines-Samuelson et al., Characterization of age-dependent and progressive cortical neuronal degeneration in presenilin conditional mutant mice. PLoS One 5, e10195 (2010).

10. K. Tabuchi, G. Chen, T. C. Südhof, J. Shen, Conditional forebrain inactivation of nicastrin causes progressive memory impairment and age-related neurodegeneration. J Neurosci 29, 7290–7301 (2009).

11. C. A. Saura et al., Loss of presenilin function causes impairments of memory and synaptic plasticity followed by age-dependent neurodegeneration. Neuron 42, 23–36 (2004).

12. H. R. Bi et al., Neuron-specific deletion of presenilin enhancer2 causes progressive astrogliosis and age-related neurodegeneration in the cortex independent of the Notch signaling. CNS Neurosci Ther 27, 174–185 (2021).

13. M. Maesako, M. C. Q. Houser, Y. Turchyna, M. S. Wolfe, O. Berezovska, Presenilin/γ-Secretase Activity Is Located in Acidic Compartments of Live Neurons. J Neurosci 42, 145–154 (2022).

14. R. Vassar et al., Beta-secretase cleavage of Alzheimer’s amyloid precursor protein by the transmembrane aspartic protease BACE. Science 286, 735–741 (1999).

15. M. Takami et al., gamma-Secretase: successive tripeptide and tetrapeptide release from the transmembrane domain of beta-carboxyl terminal fragment. J Neurosci 29, 13042–13052 (2009).

16. D. M. Bolduc, D. R. Montagna, M. C. Seghers, M. S. Wolfe, D. J. Selkoe, The amyloid-beta forming tripeptide cleavage mechanism of γ-secretase. Elife 5, (2016).

17. L. Chávez-Gutiérrez et al., The mechanism of γ-Secretase dysfunction in familial Alzheimer disease. Embo j 31, 2261–2274 (2012).

18. Y. Qi-Takahara et al., Longer forms of amyloid beta protein: implications for the mechanism of intramembrane cleavage by gamma-secretase. J Neurosci 25, 436–445 (2005).

19. S. Funamoto et al., Truncated carboxyl-terminal fragments of beta-amyloid precursor protein are processed to amyloid beta-proteins 40 and 42. Biochemistry 43, 13532–13540 (2004).

20. N. Kakuda et al., Distinct deposition of amyloid-β species in brains with Alzheimer’s disease pathology visualized with MALDI imaging mass spectrometry. Acta Neuropathol Commun 5, 73 (2017).

21. L. Fu et al., Comparison of neurotoxicity of different aggregated forms of Aβ40, Aβ42 and Aβ43 in cell cultures. J Pept Sci 23, 245–251 (2017).

22. A. J. Kuhn, J. Raskatov, Is the p3 Peptide (Aβ17-40, Aβ17-42) Relevant to the Pathology of Alzheimer’s Disease?1. J Alzheimers Dis 74, 43–53 (2020).

23. S. F. Lichtenthaler, α-secretase in Alzheimer’s disease: molecular identity, regulation and therapeutic potential. J Neurochem 116, 10–21 (2011).

24. M. D. Tambini, K. A. Norris, L. D’Adamio, Opposite changes in APP processing and human Aβ levels in rats carrying either a protective or a pathogenic APP mutation. Elife 9, (2020).

25. M. Mullan et al., A pathogenic mutation for probable Alzheimer’s disease in the APP gene at the N-terminus of beta-amyloid. Nature genetics 1, 345–347 (1992).

26. M. Pagnon de la Vega et al., The Uppsala APP deletion causes early onset autosomal dominant Alzheimer’s disease by altering APP processing and increasing amyloid β fibril formation. Science translational medicine 13, (2021).

27. I. E. Jansen et al., Genome-wide meta-analysis identifies new loci and functional pathways influencing Alzheimer’s disease risk. Nat Genet 51, 404–413 (2019).

28. S. Veugelen, T. Saito, T. C. Saido, L. Chávez-Gutiérrez, B. De Strooper, Familial Alzheimer’s Disease Mutations in Presenilin Generate Amyloidogenic Aβ Peptide Seeds. Neuron 90, 410–416 (2016).

29. M. A. Fernandez, J. A. Klutkowski, T. Freret, M. S. Wolfe, Alzheimer presenilin-1 mutations dramatically reduce trimming of long amyloid β-peptides (Aβ) by γ-secretase to increase 42-to-40-residue Aβ. J Biol Chem 289, 31043–31052 (2014).

30. M. Szaruga et al., Alzheimer’s-Causing Mutations Shift Aβ Length by Destabilizing γ-Secretase-Aβn Interactions. Cell 170, 443–456.e414 (2017).

31. S. Devkota, T. D. Williams, M. S. Wolfe, Familial Alzheimer’s disease mutations in amyloid protein precursor alter proteolysis by γ-secretase to increase amyloid β-peptides of >45 residues. J Biol Chem, 100281 (2021).

32. B. Kretner et al., Generation and deposition of Aβ43 by the virtually inactive presenilin-1 L435F mutant contradicts the presenilin loss-of-function hypothesis of Alzheimer’s disease. 8, 458–465 (2016).

33. D. Petit et al., Aβ profiles generated by Alzheimer’s disease causing PSEN1 variants determine the pathogenicity of the mutation and predict age at disease onset. Molecular Psychiatry, (2022).

34. K. G. Mawuenyega et al., Decreased clearance of CNS beta-amyloid in Alzheimer’s disease. *Science (New York*, N.Y.) 330, 1774 (2010).

35. L. Liu et al., Identification of the Aβ37/42 peptide ratio in CSF as an improved Aβ biomarker for Alzheimer’s disease. Alzheimers Dement, (2022).

36. G. Brinkmalm et al., Identification of neurotoxic cross-linked amyloid-β dimers in the Alzheimer’s brain. Brain 142, 1441–1457 (2019).

37. D. M. Walsh et al., Naturally secreted oligomers of amyloid beta protein potently inhibit hippocampal long-term potentiation in vivo. Nature 416, 535–539 (2002).

38. W. Hong et al., Diffusible, highly bioactive oligomers represent a critical minority of soluble Aβ in Alzheimer’s disease brain. Acta Neuropathol 136, 19–40 (2018).

39. Z. Wang et al., Human Brain-Derived Aβ Oligomers Bind to Synapses and Disrupt Synaptic Activity in a Manner That Requires APP. J Neurosci 37, 11947–11966 (2017).

40. I. Lauritzen, R. Pardossi-Piquard, A. Bourgeois, A. Bécot, F. Checler, Does Intraneuronal Accumulation of Carboxyl-terminal Fragments of the Amyloid Precursor Protein Trigger Early Neurotoxicity in Alzheimer’s Disease? Curr Alzheimer Res 16, 453–457 (2019).

41. W. Xu et al., Amyloid precursor protein-mediated endocytic pathway disruption induces axonal dysfunction and neurodegeneration. J Clin Invest 126, 1815–1833 (2016).

42. D. Kwart et al., A Large Panel of Isogenic APP and PSEN1 Mutant Human iPSC Neurons Reveals Shared Endosomal Abnormalities Mediated by APP β-CTFs, Not Aβ. Neuron 104, 1022 (2019).

43. S. Kim et al., Evidence that the rab5 effector APPL1 mediates APP-βCTF-induced dysfunction of endosomes in Down syndrome and Alzheimer’s disease. Mol Psychiatry 21, 707–716 (2016).

44. M. L. Franco et al., TrkA-mediated endocytosis of p75-CTF prevents cholinergic neuron death upon γ-secretase inhibition. Life Sci Alliance 4, (2021).

45. N. Matsumura et al., gamma-Secretase associated with lipid rafts: multiple interactive pathways in the stepwise processing of beta-carboxyl-terminal fragment. J Biol Chem 289, 5109–5121 (2014).

46. C. C. Shelton et al., A miniaturized 1536-well format gamma-secretase assay. Assay Drug Dev Technol 7, 461–470 (2009).

47. X. Hu et al., Amyloid seeds formed by cellular uptake, concentration, and aggregation of the amyloid-beta peptide. Proc Natl Acad Sci U S A 106, 20324–20329 (2009).

48. Y. Su, P. T. Chang, Acidic pH promotes the formation of toxic fibrils from beta-amyloid peptide. Brain Res 893, 287–291 (2001).

49. M. P. Schützmann et al., Endo-lysosomal Aβ concentration and pH trigger formation of Aβ oligomers that potently induce Tau missorting. Nat Commun 12, 4634 (2021).

50. R. Q. Liu et al., Membrane localization of beta-amyloid 1-42 in lysosomes: a possible mechanism for lysosome labilization. J Biol Chem 285, 19986–19996 (2010).

51. E. K. Esbjörner et al., Direct observations of amyloid β self-assembly in live cells provide insights into differences in the kinetics of Aβ(1-40) and Aβ(1-42) aggregation. Chem Biol 21, 732–742 (2014).

52. R. P. Friedrich et al., Mechanism of amyloid plaque formation suggests an intracellular basis of Abeta pathogenicity. Proc Natl Acad Sci U S A 107, 1942–1947 (2010).

53. M. Maesako et al., Visualization of PS/γ-Secretase Activity in Living Cells. iScience 23, 101139 (2020).

54. M. C. Houser et al., A Novel NIR-FRET Biosensor for Reporting PS/γ-Secretase Activity in Live Cells. Sensors (Basel) 20, (2020).

55. T. Schneider-Poetsch et al., Inhibition of eukaryotic translation elongation by cycloheximide and lactimidomycin. Nat Chem Biol 6, 209–217 (2010).

56. K. Uemura et al., Characterization of sequential N-cadherin cleavage by ADAM10 and PS1. Neurosci Lett 402, 278–283 (2006).

57. L. W. Wong, Z. Wang, S. R. X. Ang, S. Sajikumar, Fading memories in aging and neurodegeneration: Is p75 neurotrophin receptor a culprit? Ageing Res Rev 75, 101567 (2022).

58. J. N. Conroy, E. J. Coulson, High-affinity TrkA and p75 neurotrophin receptor complexes: a twisted affair. J Biol Chem, 101568 (2022).

59. D. M. Loeb et al., The trk proto-oncogene rescues NGF responsiveness in mutant NGF-nonresponsive PC12 cell lines. Cell 66, 961–966 (1991).

60. E. K. Pickett et al., Non-Fibrillar Oligomeric Amyloid-β within Synapses. J Alzheimers Dis 53, 787–800 (2016).

61. S. Schedin-Weiss, I. Caesar, B. Winblad, H. Blom, L. O. Tjernberg, Super-resolution microscopy reveals γ-secretase at both sides of the neuronal synapse. Acta Neuropathol Commun 4, 29 (2016).

62. P. R. Dunkley, P. E. Jarvie, P. J. Robinson, A rapid Percoll gradient procedure for preparation of synaptosomes. Nat Protoc 3, 1718–1728 (2008).

63. L. Fonseca-Ornelas et al., Altered conformation of α-synuclein drives dysfunction of synaptic vesicles in a synaptosomal model of Parkinson’s disease. Cell Rep 36, 109333 (2021).

64. P. Scheltens et al., Alzheimer’s disease. Lancet 397, 1577–1590 (2021).

65. D. S. Knopman et al., Alzheimer disease. Nat Rev Dis Primers 7, 33 (2021).

66. G. Festa et al., Aggregation States of A. Int J Mol Sci 20, (2019).

67. F. Dulin et al., P3 peptide, a truncated form of A beta devoid of synaptotoxic effect, does not assemble into soluble oligomers. FEBS Lett 582, 1865–1870 (2008).

68. L. S. Higgins, G. M. Murphy, L. S. Forno, R. Catalano, B. Cordell, P3 beta-amyloid peptide has a unique and potentially pathogenic immunohistochemical profile in Alzheimer’s disease brain. Am J Pathol 149, 585–596 (1996).

69. M. López de la Paz, L. Serrano, Sequence determinants of amyloid fibril formation. Proc Natl Acad Sci U S A 101, 87–92 (2004).

70. A. J. Kuhn, B. S. Abrams, S. Knowlton, J. A. Raskatov, Alzheimer’s Disease "Non-amyloidogenic" p3 Peptide Revisited: A Case for Amyloid-α. ACS Chem Neurosci 11, 1539–1544 (2020).

71. T. C. Saido, W. Yamao-Harigaya, T. Iwatsubo, S. Kawashima, Amino- and carboxyl-terminal heterogeneity of beta-amyloid peptides deposited in human brain. Neurosci Lett 215, 173–176 (1996).

72. M. Lalowski et al., The "nonamyloidogenic" p3 fragment (amyloid beta17-42) is a major constituent of Down’s syndrome cerebellar preamyloid. J Biol Chem 271, 33623–33631 (1996).

73. E. Gowing et al., Chemical characterization of A beta 17-42 peptide, a component of diffuse amyloid deposits of Alzheimer disease. J Biol Chem 269, 10987–10990 (1994).

74. D. M. Vadukul et al., Internalisation and toxicity of amyloid-β 1-42 are influenced by its conformation and assembly state rather than size. FEBS Lett 594, 3490–3503 (2020).

75. E. Wesén, G. D. M. Jeffries, M. Matson Dzebo, E. K. Esbjörner, Endocytic uptake of monomeric amyloid-β peptides is clathrin- and dynamin-independent and results in selective accumulation of Aβ(1-42) compared to Aβ(1-40). Sci Rep 7, 2021 (2017).

76. D. Ling, M. Magallanes, P. M. Salvaterra, Accumulation of amyloid-like Aβ1-42 in AEL (autophagy-endosomal-lysosomal) vesicles: potential implications for plaque biogenesis. ASN Neuro 6, (2014).

77. B. A. Bahr et al., Amyloid beta protein is internalized selectively by hippocampal field CA1 and causes neurons to accumulate amyloidogenic carboxyterminal fragments of the amyloid precursor protein. J Comp Neurol 397, 139–147 (1998).

78. A. Sotthibundhu et al., Beta-amyloid(1-42) induces neuronal death through the p75 neurotrophin receptor. J Neurosci 28, 3941–3946 (2008).

79. E. J. Coulson et al., p75 neurotrophin receptor mediates neuronal cell death by activating GIRK channels through phosphatidylinositol 4,5-bisphosphate. J Neurosci 28, 315–324 (2008).

80. A. M. Weissmiller et al., A γ-secretase inhibitor, but not a γ-secretase modulator, induced defects in BDNF axonal trafficking and signaling: evidence for a role for APP. PLoS One 10, e0118379 (2015).

81. M. Sawa et al., Impact of increased APP gene dose in Down syndrome and the Dp16 mouse model. Alzheimers Dement 18, 1203–1234 (2022).

82. A. Salehi et al., Increased App expression in a mouse model of Down’s syndrome disrupts NGF transport and causes cholinergic neuron degeneration. Neuron 51, 29–42 (2006).

83. Y. Jiang et al., Lysosomal Dysfunction in Down Syndrome Is APP-Dependent and Mediated by APP-βCTF (C99). J Neurosci 39, 5255–5268 (2019).

84. P. Hou et al., The γ-secretase substrate proteome and its role in cell signaling regulation. Mol Cell 83, 4106–4122.e4110 (2023).

85. M. Koch et al., APP substrate ectodomain defines amyloid-β peptide length by restraining γ-secretase processivity and facilitating product release. EMBO J 42, e114372 (2023).

86. J. McInnes et al., Synaptogyrin-3 Mediates Presynaptic Dysfunction Induced by Tau. Neuron 97, 823–835.e828 (2018).

87. L. M. Ittner et al., Dendritic function of tau mediates amyloid-beta toxicity in Alzheimer’s disease mouse models. Cell 142, 387–397 (2010).

88. E. D. Roberson et al., Reducing endogenous tau ameliorates amyloid beta-induced deficits in an Alzheimer’s disease mouse model. Science 316, 750–754 (2007).

89. T. L. Spires-Jones, B. T. Hyman, The intersection of amyloid beta and tau at synapses in Alzheimer’s disease. Neuron 82, 756–771 (2014).

90. P. Ferrer-Raventós et al., Amyloid precursor protein. Neuropathol Appl Neurobiol 49, e12879 (2023).

91. M. Pera et al., Distinct patterns of APP processing in the CNS in autosomal-dominant and sporadic Alzheimer disease. Acta Neuropathol 125, 201–213 (2013).

92. M. Szaruga et al., Qualitative changes in human γ-secretase underlie familial Alzheimer’s disease. J Exp Med 212, 2003–2013 (2015).

93. J. Shen, R. J. Kelleher, The presenilin hypothesis of Alzheimer’s disease: evidence for a loss-of-function pathogenic mechanism. Proc Natl Acad Sci U S A 104, 403–409 (2007).

94. M. Maesako et al., Pathogenic PS1 phosphorylation at Ser367. Elife 6, (2017).

95. L. Chavez-Gutierrez et al., The mechanism of gamma-Secretase dysfunction in familial Alzheimer disease. EMBO J 31, 2261–2274 (2012).

96. A. Gore et al., Somatic coding mutations in human induced pluripotent stem cells. Nature 471, 63–67 (2011).

97. J. E. Young et al., Elucidating molecular phenotypes caused by the SORL1 Alzheimer’s disease genetic risk factor using human induced pluripotent stem cells. Cell Stem Cell 16, 373–385 (2015).

98. L. K. Fong et al., Full-length amyloid precursor protein regulates lipoprotein metabolism and amyloid-β clearance in human astrocytes. J Biol Chem 293, 11341–11357 (2018).

99. U. Das et al., Activity-induced convergence of APP and BACE-1 in acidic microdomains via an endocytosis-dependent pathway. Neuron 79, 447–460 (2013).

100. G. M. Shankar, A. T. Welzel, J. M. McDonald, D. J. Selkoe, D. M. Walsh, Isolation of low-n amyloid β-protein oligomers from cultured cells, CSF, and brain. Methods Mol Biol 670, 33–44 (2011).

101. K. Johnson-Wood et al., Amyloid precursor protein processing and A beta42 deposition in a transgenic mouse model of Alzheimer disease. Proc Natl Acad Sci U S A 94, 1550–1555 (1997).

102. H. Wickham, in Use R!,. (Springer International Publishing : Imprint: Springer,, Cham, 2016), pp. 1 online resource (XVI, 260 pages 232 illustrations, 140 illustrations in color.

103. H. Wickham. (Journal of Statistical Software, 2007), vol. 21, pp. 1–20.

104. H. Wickham. (Journal of Statistical Software, 2011), vol. 40, pp. 1–29.

